# Scaling Multiplex qPCR Primer Design to 1000-plex using the Degenerate Incomplete Multiplex Primer List Extension (DIMPLE) Algorithm

**DOI:** 10.64898/2026.04.17.719221

**Authors:** Alessandro Pinto, Xue Dong, Wei Wu, Sara J. Johnson, Qin Wen, Cheng Zhang, Jana Havey, Bonnie Wang, Guilin Tang, Aziz Farhat, David Yu Zhang, Ghayas C. Issa, Xi Zhang

## Abstract

Massively multiplexed qPCR is primarily constrained by increasing primer dimer formation as the number of distinct primers in a single reaction increases. Previous multiplex primer design algorithms either fail to sufficiently suppress primer dimers at 100+ plex, or take exceedingly high amounts of computational resources to complete. Here, we present DIMPLE, a linear-runtime primer design algorithm that effectively generates 10,000+ primers to amplify thousands of potential amplicons in a single qPCR reaction. As one clinical demonstration of this algorithm, we designed an assay to detect 2,302 distinct KMT2A gene fusion subtypes using 204 primers in a single tube. In contrast to FISH and convention NGS approaches with 2% variant allele frequency (VAF) limit of detection, our DIMPLE qPCR assay was able to analytically detect gene fusions down to 0.05% VAF. We also constructed proof-of-concept multiplex qPCR panels for additional oncology gene fusions, multiplex pathogen detection, and DNA methylation markers. The scalability and low computational cost DIMPLE are complementary to new instrument platforms for massively multiplex qPCR readout for enabling rapid, point-of-care nucleic acid testing.

## Introduction

Polymerase chain reaction (PCR) is the gold standard today for molecular diagnostics based on detection of nucleic acids, particularly in clinical in vitro diagnostics (IVD) settings^1,2^. However, because the number of clinically important nucleic acid targets is dramatically rising due to high-throughput discovery studies based on next-generation sequencing (NGS), current standard qPCR instruments with 4 to 6 spectrally distinct fluorescent channels are no longer sufficient for sensitive detection of many clinically validated nucleic acid targets^3^. Simultaneously, NGS wet lab and bioinformatics processes remains slow, complex, and expensive^4^. The combination of *multiplex limitations* of qPCR and the *complexity limitations* of NGS has created a need for highly multiplex qPCR for in-hospital diagnostics, particularly for point-of-care testing (PoCT) applications when medical conditions are acute^5^. A number of massively multiplex qPCR instrument platforms (Biofire^6^, Torus^7^, Flash Dx^8^, Cepheid^5^, QiaSTAT^9^) have been commercially introduced in the last few years to enable higher plex qPCR, but these systems solve the readout problem and not the multiplex PCR amplification problem.

Primer dimers are artifacts formed when two PCR primers unintentionally bind to each other and extend off each other^1,10^. Primer dimers are the primary culprit for the failure of massively qPCR reactions, because primer dimers deplete the primers intended for amplification of target nucleic acid sequences, resulting in false negatives in Taqman-based assays and false positives in intercalating dye (e.g. SYBR) assays^11^. The number of possible primer dimers increase quadratically as the number of distinct primers in a single reaction, with N*(N+1)/2 possible primer dimer species for a total of N primers in a single reaction^11,12^. This makes scaling of multiplex primers far more difficult than may be expected intuitively. For example, a primer design algorithm that produces 2 functional primers with 99% success rate would have less than a 1 in 20 million chance of successfully design a set of 200 primers (0.99^(5050/3)). In NGS settings, primer dimers can be removed via size selection using beads or columns,^13^ but these solid-phase purification methods are not compatible with sample-in, answer-out qPCR instruments and assays.

Computational algorithms for massively multiplex PCR primer design, such as the SADDLE algorithm^11^ and related methods^14–16^, generally depend on two tools **(1)** heuristics to estimate the likelihood of primer dimer formation between 2 primers, in order to estimate ensemble primer dimer formation from N primers, and **(2)** a non-convex optimization algorithm for replacing individual primers with alternative primer candidates for amplifying the same DNA target sequence. The problem with previous approaches is that **(2)** becomes inordinately computationally expensive as N increases, because the time complexity of SADDLE and other algorithms are at least O(N^2)^12^, even disregarding nonlinearities in parameter space exploration. Even with modern high-throughput computing infrastructure, primer set design can take weeks to months for N≥10,000. Worse yet, these multiplex primer design algorithms can be trapped in highly suboptimal primer sets^17^ when one or more DNA target exhibit pathological properties (e.g. highly repetitive sequence) that effectively prevent primer design. Finally, **(1)** is imperfect, and some amount of empirical optimization is needed to replace the primers that experimentally contributes to primer dimers which are not predicted by primer dimer evaluation heuristics^18,19^.

Here, we present the Degenerate Incomplete Multiplex Primer List Expansion (DIMPLE) algorithm for massively multiplex PCR primer design. At its core, DIMPLE starts with an empty primer set and then adds primers from a primer candidate pool using a linear time greedy algorithm. This is in contrast to starting from a randomly generated complete set of primers^17^, and using a quadratic time simulated annealing algorithm to replace primers piecemeal (Fig. 1a). Due to the time complexity advantage of this approach, we typically run DIMPLE to design a primer set that is 5x redundant, meaning for every single target DNA sequence, we design up to 5 distinct forward primers and 5 distinct reverse primers. This allows us significant flexibility in the experimental portion of optimizing a high-plex panel, because we essentially replace primers from a pre-validated pool of alternative primers that are designed to be compatible with the remaining primers.

**Fig. 1:**
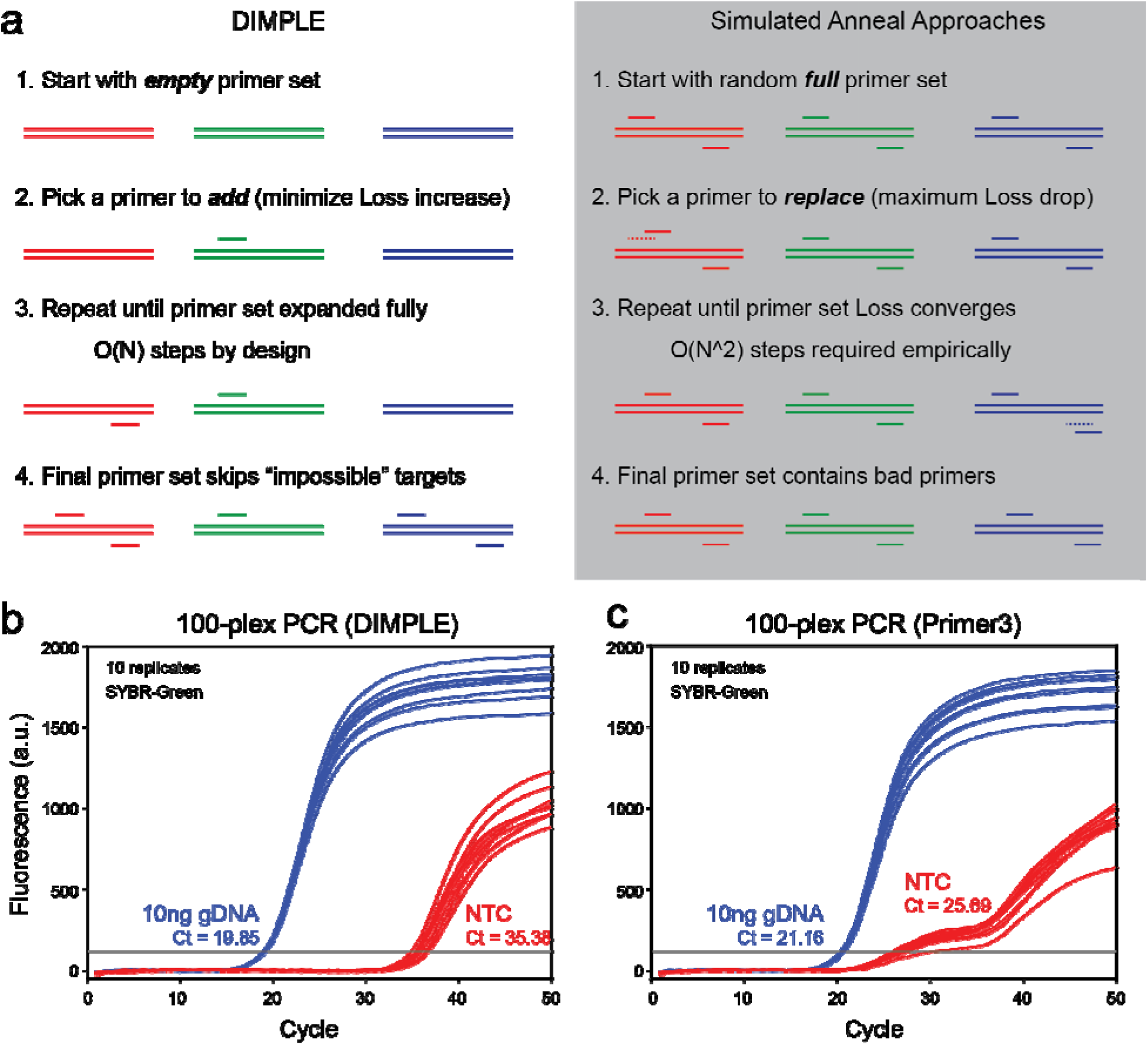
DIMPLE algorithm for designing highly multiplex PCR primer sets. **(a)** Schematic of DIMPLE vs. SADDLE and other simulated annealing-based primer design approaches. **(b)** First-pass design of a 100-plex (200 primer) qPCR reaction using DIMPLE. SYBR-Green intercalating use was used to visualize the relative amplification of desired amplicons vs. primer dimers in the no-template control (NTC) reaction. The qPCR trace using 10ng human gDNA (3000 haploid copies) shows ΔCt > 15 vs. the NTC. **(c)** First-pass 100-plex qPCR primer design by the Primer3 software shows ΔCt < 5, and NTC Ct = 25.7, indicating significant primer dimer formation.

## Results

### 100-plex Panel Comparison vs. Primer 3

We begin by evaluating DIMPLE vs. Primer3^20^ on a 200 primer (100-plex) qPCR assay without any empirical optimization (Fig. 1b and 1c). Here, the 100 target sequences are randomly chosen regions of the human genome, and the fluorescence was observed using the SYBR-Green intercalating dye. As expected, the primer set designed by DIMPLE exhibited a qPCR cycle threshold (Ct) value of 35.4 cycles for the No Template Control (NTC) reaction, significantly better than the Primer3-designed panel with NTC Ct of 25.7. For the positive control (10ng human genomic DNA), the DIMPLE panel’s Ct is 19.9 and lower than the Primer3 panel’s Ct of 21.1, indicating higher amplification yield for DIMPLE in addition to improved primer dimer suppression.

### Design Success Evaluation and Scaling to 10,000+ Primers

We next used DIMPLE to design primers for simultaneously amplifying 1000 DNA target loci associated with pathogenic mutations in 364 genes observed in lung, colorectal, breast, ovarian, and pancreatic cancers^21^. To ensure robustness of the final panel, we set DIMPLE to redundantly design 7 distinct forward primers and 7 distinct reverse primers for <u>each</u> amplicon of interest, totaling 14,000 primers. Due to the incompleteness nature of the DIMPLE algorithm, we ended up generating 11,627 unique primers (*Supplementary Data 1*) that collectively cover each DNA target region with a minimum of 3 forward and 3 reverse primers each (Fig. 2a). From this primer pool, we first arbitrarily selected 1 forward primer and 1 reverse primer per locus, for a total of 2,000 primers for first-pass experimental testing.

**Fig. 2:**
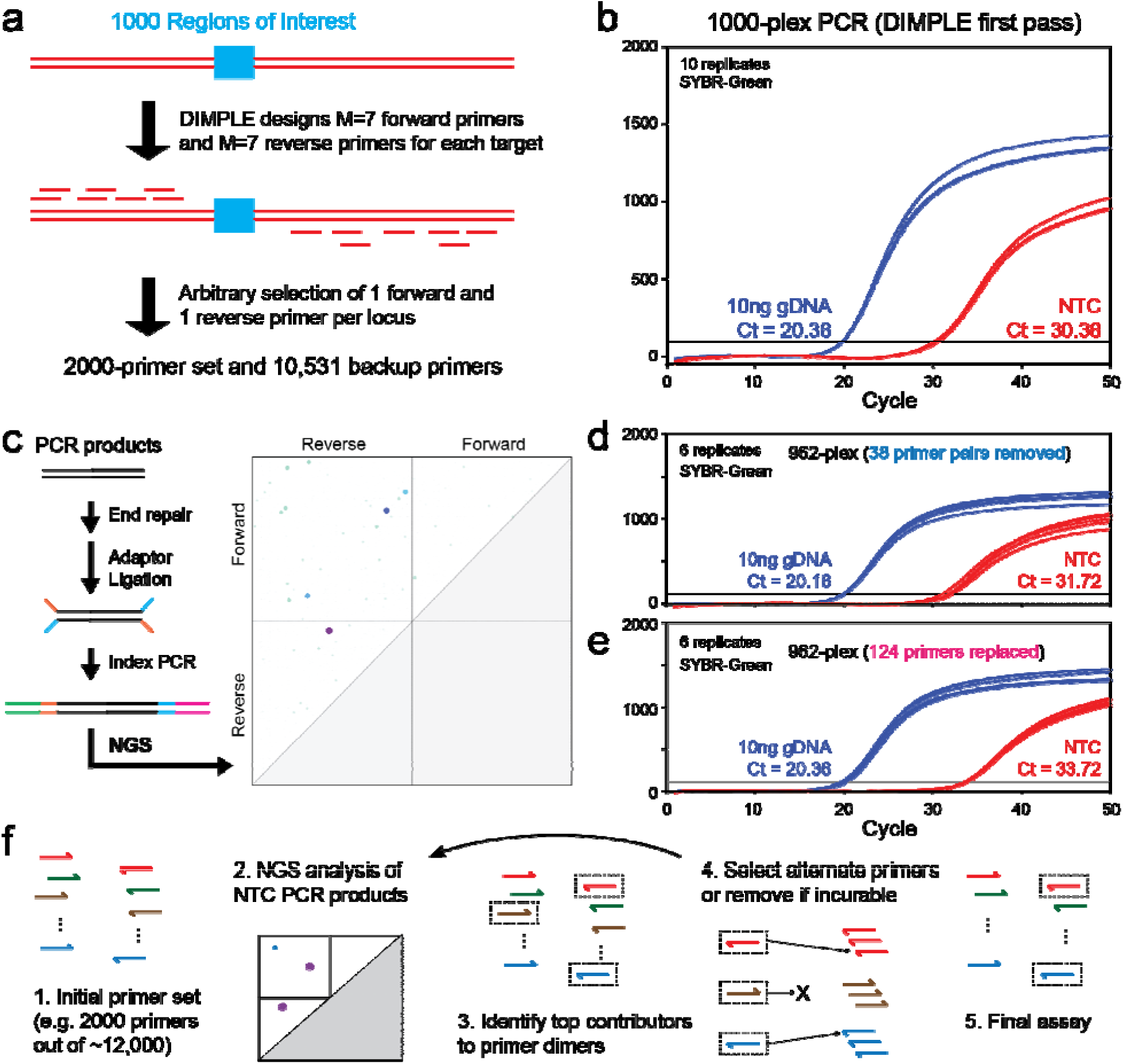
Degeneracy of primer design to facilitate experimental panel optimization. **(a)** Degenerate primer design and selection schematic. **(b)** First-pass 1000-plex DIMPLE-designed qPCR reaction; see Supplementary Section S1 for panel content details. Due to the 10-fold higher number of distinct primers, the panel’s ΔCt = 10.0 was lower than for the 100-plex panel earlier. **(c)** NGS-based analysis of NTC PCR products reveals primary contributors to primer dimer formation. **(d)** Removal of 38 primer pairs that dominate primer dimers increases the ΔCt to 11.5. **(e)** Replacing primers that contribute to primer dimer formation with alternatives from the pre-designed primer pool. Replacing 124 primers further improves the ΔCt to 13.4. **(f)** Schematic for overall process of experimental optimization of high-plex panels using redundantly designed PCR primers.

We first tested the full 1000-plex qPCR assay in a single SYBR-Green qPCR reaction, and observed a positive control Ct value of 20.4 using 10ng of human genomic DNA, corresponding to 3,000 haploid genome copies (Fig. 2b). The NTC Ct of this reaction was 30.38, indicating that human genomic DNA could be detectable using this assay down to roughly 10pg, or 3 haploid genomic copies. Although we consider this to be a very good initial result in the absence of any optimization, we want to show that significant improvements are possible through replacement of poor-performing primers.

Analysis of the NTC PCR amplification products using NGS identified the primary contributors of primer dimers. As expected, a small number of primers contributed disproportionately to the total primer dimers (Fig. 2c). We also analyzed the amount of nonspecific genomic amplification attributed to each primer. We chose to remove 38 problematic primer pairs (3.8%) with both high primer dimer contribution and high nonspecific genomic amplification. Based on NGS analysis, this would reduce the total primer dimer quantity by roughly 60%. The experimental qPCR NTC value of 31.72 for the resulting 962-plex qPCR panel (1924 primers) was 1.3 cycles delayed compared to the NTC of the original 1000-plex qPCR panel, confirming the reduced primer dimer formation (2^(-1.3) ≈ 0.4).

Next, we identified primers suitable for replacement by alternative primers from the predesigned pool to further reduce primer dimer formation. A total of 124 primer pairs were selected for replacement, and the experimental qPCR NTC value for the revised 962-plex panel was 33.72 (Fig. 2d). Compared to the original 1000-plex panel’s Ct value of 30.38, this corresponds to a total of 90% primer dimer suppression, and ΔCt value of 13.36 relative to the positive control (10ng genomic DNA) indicates an equivalent primer dimer amount as 10ng * 2^(-13.36) = 0.95pg of gDNA input, or 0.29 haploid genome equivalents. **<u>This indicates that the revised 962-plex panel is already sensitive to single genomic copy inputs</u>**, so we did not attempt further rounds of primer replacement to additionally suppress the Ct value of NTC. In a Taqman probe-based qPCR assay, such NTC amplification of primer dimers would be invisible and would not interfere with the signal from single-digit molecules of input. A summary of the overall primer set design and optimization process is shown in Fig. 2e.

### Detection of 2,302 KTM2A Gene Fusions with a 204-primer qPCR Assay

In acute myeloid leukemia (AML), roughly 5% of adults patients and 15% of pediatric patients have a rearrangement of the *KMT2A* gene (*KMT2Ar*)^22^. *KMT2A fusions* occur with up to 130 fusion gene partners and numerous potential breakpoints which has limited use of this founding genetic event for disease monitoring except for development of bespoke, patient-specific probes^23^. In addition, the advent of menin inhibitors as a targeted therapy for *KMT2Ar* leukemia, there is an unmet need for identifying the presence of these genetic rearrangements with high sensitivity and specificity, with prognostic and predictive implications informing disease recurrence monitoring and the choice of treatment. The menin inhibitor revumenib was recently approved by the Food and Drug Administration for relapsed or refractory acute leukemia with *KMT2Ar* in adult and pediatric patients 1 year and older^24^. Additionally, multiple other menin inhibitors are currently in clinical development^25^. These *KMT2Ar+* patients historically have had very poor prognosis when treated with conventional chemotherapy, however, menin inhibitors are showing promising efficacy as monotherapy or in combination with other agents for this leukemia substet^26–28^. Detection of KMT2A gene fusions would thus be informative to clinicians treating AML patients, as treatment with menin inhibitors can significantly improve patient outcomes. The technical challenge is the *KMT2A* forms fusions with numerous genes at many different breakpoints, with over 2000 known gene fusion subtypes^23^. This makes *KMT2A* gene fusion detection previously impossible for qPCR, and clinicians instead rely on either break-apart fluorescent in situ hybridization (FISH) or NGS^29^. However, not only are FISH and NGS expensive and slow, their sensitivity is limited to about 2% VAF for *KMT2A* fusions^27,30^. The VAF of *KMT2A* gene fusions in peripheral blood mononuclear cells (PBMCs) from patients may be significantly lower than 2%^31^.

We designed a 204 primer single-tube qPCR panel that detects the 2,302 gene fusion subtypes for *KMT2Ar* (Fig. 3a), covering 95%+ of reported *KMT2A* gene fusion cases^23^. The panel consists of 12 forward primers to each of the 12 exons of *KMT2A* involved in gene fusions, and 192 reverse primers to exons from the 12 most observed fusion partner genes. For this assay, **<u>to prevent false positive signal from primer dimers, we used 12 Taqman</u> <u>probes downstream of each *KMT2A* forward primer</u>**. The assay was applied to the cDNA generated from reverse transcription of RNA from reference and clinical samples.

**Fig. 3:**
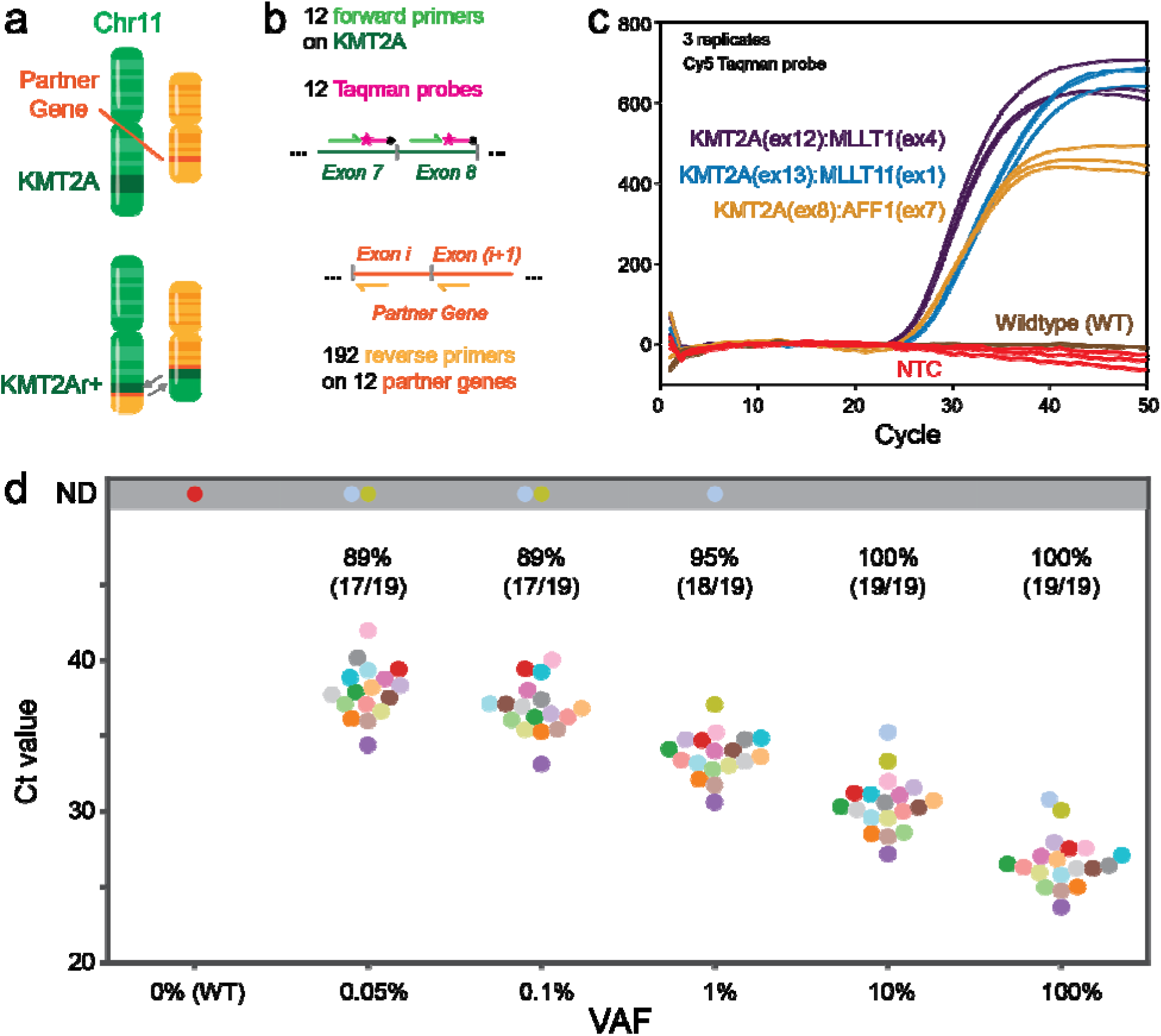
Using DIMPLE to sensitively and affordably detect oncology gene fusions. **(a)** The KMT2A gene resides on human chromosome 11, and forms gene fusions in approximately 5% of acute myeloid leukemia (AML) patients. **(b)** Design of a 204-primer panel to detect 2,304 KMT2A gene fusion subtypes that collectively span 95%+ of known KMT2A fusions. Here, to prevent false positives for clinical samples, we use Cy5-labeled Taqman probes for readout, rather than SYBR-Green intercalating dye, so that primer dimers do not generate signal. **(c)** Sample qPCR results for 3 synthetic DNA targets with different gene fusion identities. **(d)** Assessing the lower limit of detection for gene fusions using the qPCR panel. The qPCR assay is able to detect 17 out of the 19 the gene fusion subtypes tested at 0.05% VAF. We believe that this represents a fair assessment of our panel’s sensitivity to the 2,304 gene fusion subtypes, as we did not specifically optimize the panel against these 19 species.

To analytically validate the assay, we selected 19 representative *KMT2A* targets as positive controls. Of these, 5 were high-frequency fusions from FusionGDB^32^ (*KMT2A::AFF1* (exon 8, 7), *KMT2A::MLLT3* (exon 8, 6), *KMT2A::MLLT3* (exon 9, 2), *KMT2A::EPS15* (exon 10, 2), and *KMT2A::USP2* (exon 22, 3). The remaining constructs were chosen randomly to ensure at least one design per *KMT2A* exon. Each construct was ordered from commercial vendors as a 500nt double-stranded DNA fragment consisting of 250nt upstream and 250nt downstream of the breakpoint. All 19 *KMT2A* targets were clearly detected with no confounding signal from the wildtype negative control or the NTC negative control (Fig. 3b, Supp Fig. S1).

Next, we performed a titration series to characterize the lower limit of detection of the panel. In each experiment, one of the synthetic *KMT2A* fusion targets was blended at known concentration into a matrix of cDNA reverse transcribed from a consented healthy donor. The healthy donor’s *KMT2A* RNA expression level was characterized by qPCR, and then used to generate reference samples with KMT2A gene fusion variant allele frequencies (VAF) between 100% and 0.05%. This blending experiment was performed for all 19 targets at each of the 5 VAF values tested. In 17 out of 19 cases, the qPCR assay successfully detects the RNA fusion at 0.05% VAF.

### Application of KTM2A Gene Fusion qPCR Panel to Clinical Samples

We next applied the 204-primer KMT2A qPCR assay to 16 blinded retrospective AML samples collected at MD Anderson Cancer Center (Fig. 4a). The assay demonstrated high specificity, correctly identifying samples with non-actionable *KMT2A* rearrangement as negative, and resolved ambiguous cases where FISH was inconclusive or unavailable. One sample (MD15) harboring a rare t(2;11) translocation (*KMT2A::THADA*) tested negative, as expected given that THADA is not among the 12 fusion partners covered by the panel. Demographic and cytogenetic characteristics are provided in *Supplementary Tables S2–S4*.

**Fig. 4:**
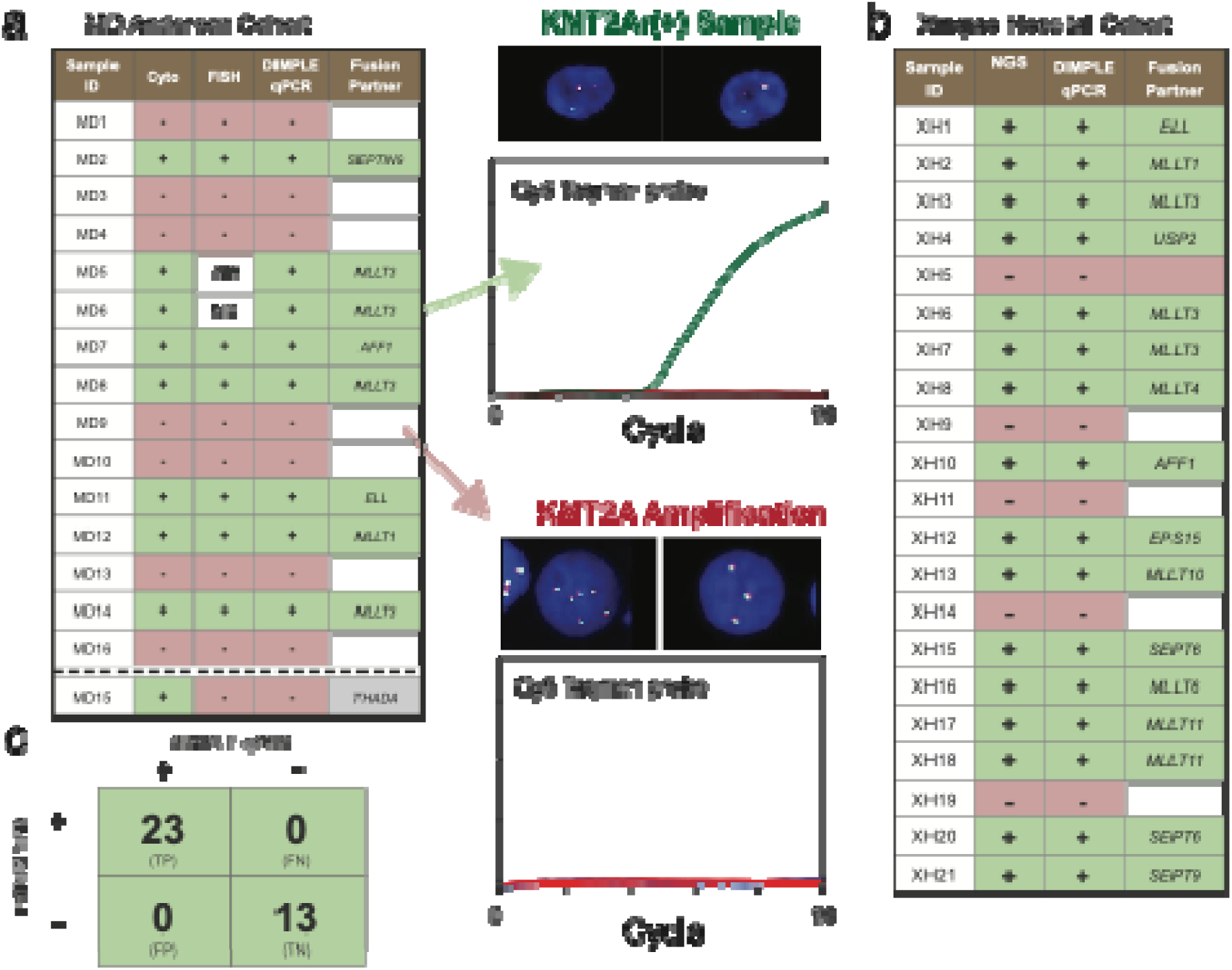
Validation of the KMT2A gene fusion qPCR panel vs. retrospective clinical samples. **(a)** Concordance between cytogenetics, FISH, and DIMPLE qPCR across 16 blinded AML samples from MD Anderson Cancer Center. Sample MD15 (dashed) harbors a rare KMT2A::THADA fusion not included in the panel. ND, not done. Center: representative FISH images and qPCR curves for a KMT2Ar-positive sample (green) versus KMT2A amplification without fusion (red). **(b)** Independent validation at Xinqiao Hospital comparing DIMPLE qPCR to NGS across 21 samples. **(c)** Combined confusion matrix for both cohorts.

To evaluate assay transferability, we shipped a kit to Xinqiao Hospital (Chongqing, China), where 21 retrospective samples were independently tested. The assay achieved 100% concordance with NGS, correctly identifying all 16 *KMT2Ar*-positive and 5 *KMT2Ar*-negative samples (Fig. 4b). Across both cohorts, the assay demonstrated 100% sensitivity (23/23) and 100% specificity (13/13) for covered fusion targets (Fig. 4c).

Given the improved VAF limit of detection of our qPCR assay, we expect that if we test sufficient additional clinical samples that were assessed as *KMT2Ar-* by break apart FISH, we would identify some amount of false negatives, in which *KMT2A* gene fusions were present but at VAFs lower than the 2% detectable limit by FISH.

### Other Oncology Gene Fusion Panels

Using the same approach as for *KMT2A* gene fusions, we also designed a 99-primer NTRK fusions qPCR panel, and a 130-primer *EWSR1+FUS* fusions qPCR panel. The NTRK panel consists of 19 primers on the *NTRK1*, *NTRK2*, and *NTRK3* genes, and 80 primers across 53 most common partner genes. Importantly, the FDA approved NTRK inhibitors Entrectinib^33^ (Roche/Ignyta), Larotrectinib (Lilly/Loxo), and Repotrectinib (BMS)^34^ are approved for **<u>all solid tumors</u>** that exhibit an *NTRK* gene fusion^35^. Heretofore, the challenge has been identifying these *NTRK* fusion patients, because they are generally less than 1% of the overall solid tumor patient population, except in select cancer types such as sarcomas where NTRK fusions can be highly enriched. On contrived and reference samples, the NTRK fusion panel was able to detect down to single digit copies of the target gene fusions.

Two other genes with frequent fusions in sarcomas are *EWSR1* and *FUS*. Like *KMT2A* and *NTRK1/2/3*, the *EWSR1* and *FUS* genes each can fuse promiscuously with many different gene partners^36^. However, unlike *KMT2A* and *NTRK1/2/3*, there are not targeted therapies directly positively indicated by the detection of *EWSR1* or *FUS* gene fusions in patient samples. However, clinicians still generally want information on *EWSR1* and *FUS* fusion status, both because they are diagnostically and prognostically important^37^ and because a number of drug candidates are currently in clinical trials (e.g. BET inhibitors^38^ and KDM1A inhibitors^39^). Our 130-primer panel includes 14 primers for *EWSR1*, 12 primers for FUS, and 104 primers across 10 high-frequency partner genes (*ATF1*, *CREB1*, *ERG*, *ETV1*, *ETV4*, *FEV*, *FLI1*, *NFATc2*, *POU5F1*, and *SMARCA5*). On contrived and reference samples, the *EWSR1+FUS* fusion panel was able to detect down to single digit copies of the target gene fusions.

### Proof of Concept for Methylation Marker Detection

To demonstrate the generality of the DIMPLE approach, we next used DIMPLE to design primer sets for detection of methylation markers. A significant portion of cytosines directly upstream of guanines (CpG sites) have the 5methylcytosine (5mC) epigenetic modification. The presence of a high concentration of 5mCs in the promoter region of a gene is known to reduce transcriptional expression of the gene^40,41^. Importantly, different organs and tissues in the human body exhibit different DNA methylation patterns, so tissue-of-origin analyses for circulating tumor cells and circulating tumor DNA are typically based on DNA methylation analysis^42,43^. Given the high chemical similarity of 5mC and unmethylated cytosines, DNA typically needs to be either chemically treated with bisulfite conversion^44^ or enzymatically treated with APOBEC^45^. Through either process, unmethylated cytosines are converted to uracils, while 5mC remain as 5mC nucleotides.

The additional design challenge of advantage of designing massively multiplex PCR primers for bisulfite converted DNA for methylation profiling is that the converted DNA has (1) low sequence complexity with very few C’s remaining, (2) low G/C content, and (3) distinct noncomplementary target sequences resulting from the conversion of the + and - strands. Designing highly multiplex PCR primers for profiling methylation markers is thus generally more difficult than standard multiplex PCR primer design. Here, we design a proof-of-concept panel covering 20 DNA loci, comprising 80 primers (40 primer pairs for converted + strand targets, 40 primer pairs for converted - strand targets) and using SYBR-Green intercalating dye to assess degree of primer dimer formation (Fig. 6). By default, DIMPLE minimizes CpG dinucleotides near primer 3′ ends, ensuring unbiased amplification of both methylated and unmethylated templates. Additionally, because bisulfite conversion renders the two DNA strands non-complementary, DIMPLE designs primer pairs for both converted strands, enabling simultaneous amplification in a single reaction.

**Fig. 5:**
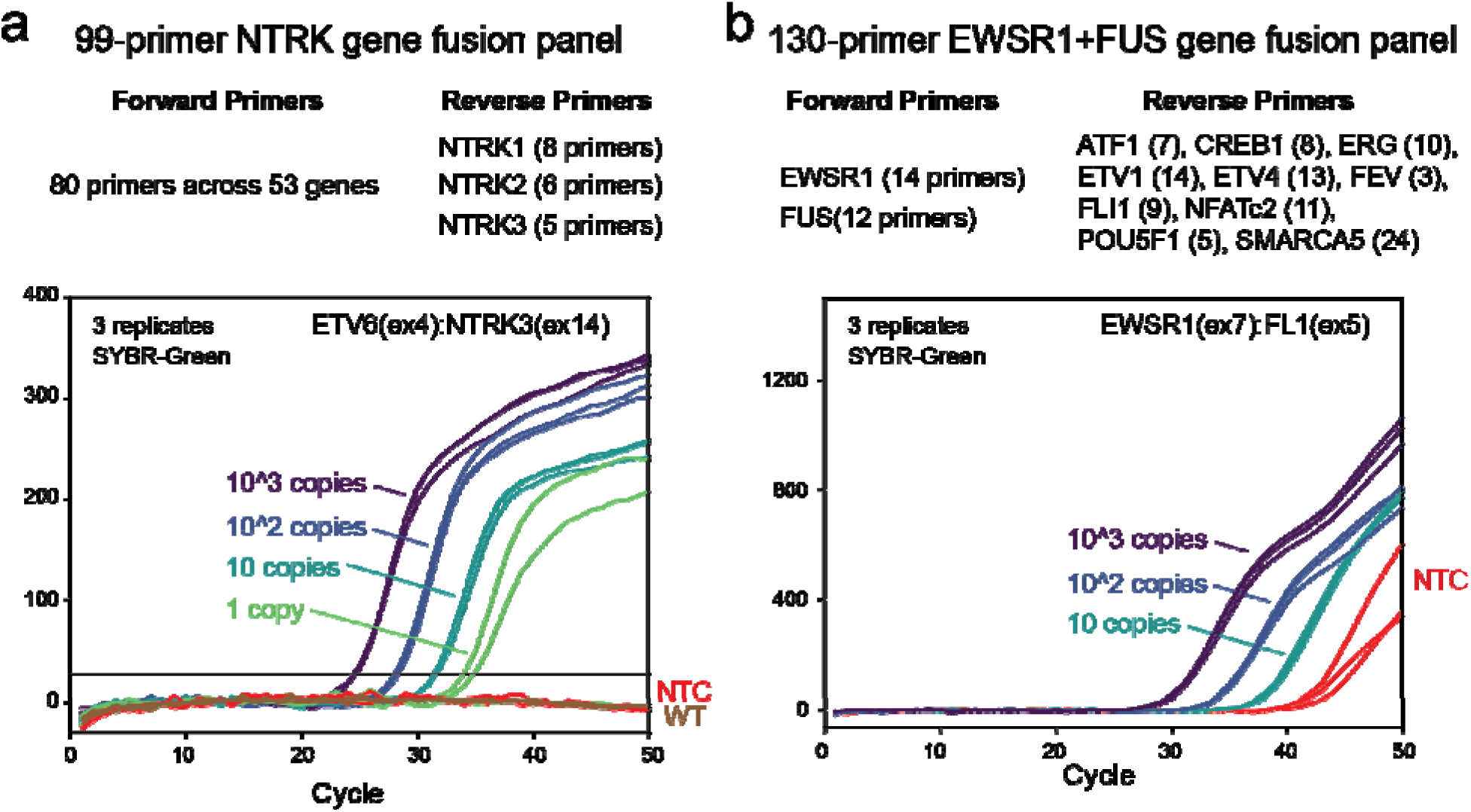
Two additional oncology gene fusion qPCR panels. **(a)** NTRK gene fusion panel detecting 1,520 gene fusion subtypes commonly observed in sarcomas; see Supplementary Section S3 for panel content details. Despite being a SYBR-Green based qPCR assay, this assay performed remarkably well with no observable signal for either the NTC or the Wildtype (WT) inputs. **(b)** EWSR1 and FUS gene fusion panel detecting 2,704 gene fusion subtypes commonly observed in sarcomas. Here, 10 gene fusion copies were detectable vs. the NTC control.

**Fig. 6:**
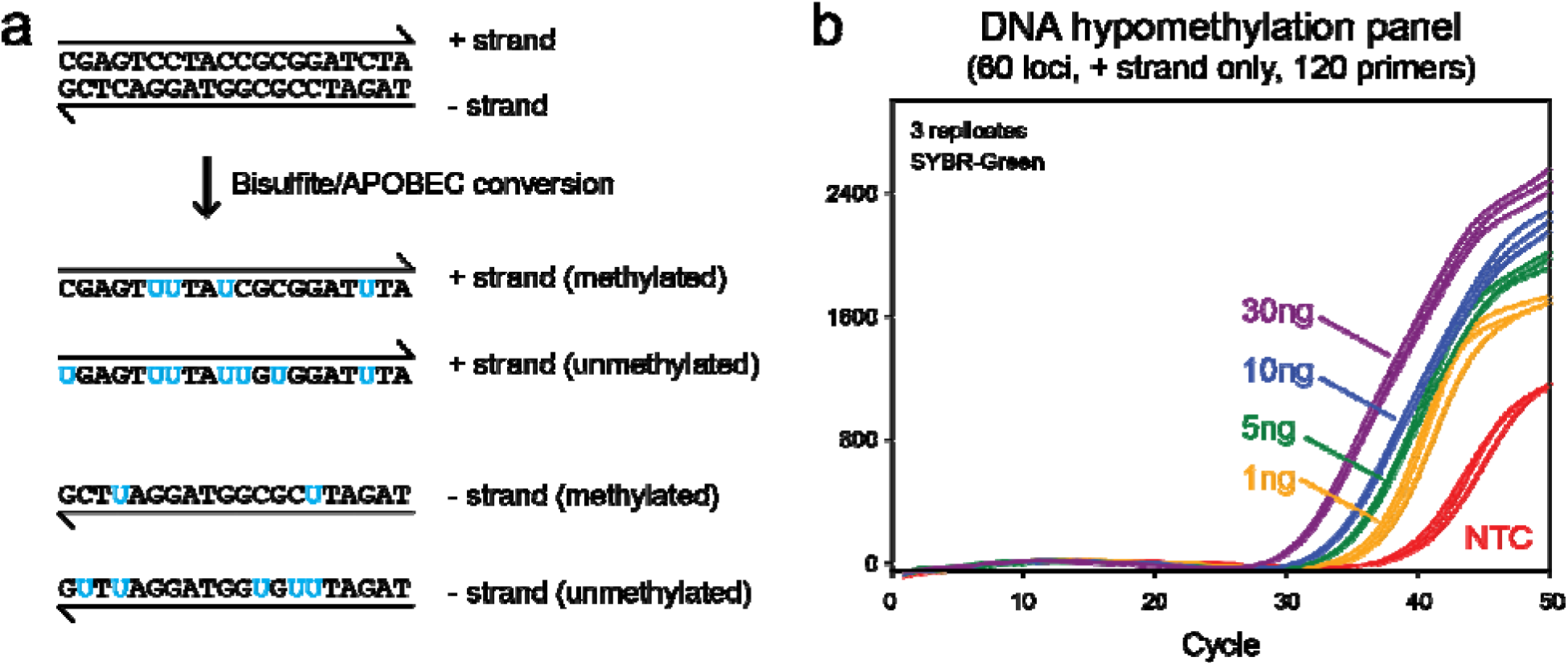
Demonstration of DIMPLE multiple qPCR primer design for DNA methylation marker detection. **(a)** Illustration of the products of bisulfite/APOBEC conversion of DNA. For each DNA locus, up to 4 different sets of primers (8 primers) must be designed to maximize sensitivity to the insert sequence. Alternatively, primer specificity can be used to detect specific methylation states of the primer binding regions. **(b)** Demonstration of 60-plex qPCR amplification of DNA hypomethylation markers. Here, 120 primers were designed to amplify only the + strands of the converted unmethylated DNA molecules, because unmethylated DNA has the lower sequence diversity than converted methylated DNA molecules, and is more challenging for traditional PCR primer design algorithms.

Fully methylated and fully unmethylated control DNA (10 ng and 1 ng inputs) were subjected to bisulfite conversion prior to amplification. Both methylation states amplified with near-equivalent efficiency: at 10 ng input (∼30.3 cycles), ΔCq was <1 cycle; at 1 ng input, ΔCq was ∼1 cycle (34.3 vs 35.3 for unmethylated vs methylated). Cq values were delayed approximately 10 cycles relative to unconverted DNA, reflecting template degradation inherent to bisulfite treatment. Nevertheless, the DIMPLE-designed primer set effectively suppressed primer dimer artifacts, and even 1 ng bisulfite-converted input was clearly distinguishable from the no-template control.

## Discussion

Massively multiplexed qPCR at the point of care is necessary for improving patient outcomes for a variety of medical diseases. Increasing the number of nucleic acid targets detectable and quantifiable in a single reaction results in 3 desirable features: (1) lowered costs from lowered number of reaction consumables, (2) lowered sample input volume requirements, and (3) improved sensitivity to via detection of increased numbers of distinct nucleic acid biomarkers. In infectious diseases, bacteria and viruses are constantly mutating under selective pressure from antimicrobials and antivirals^46^; in oncology, tumor cells are constantly mutating under selective pressure from the host immune system and targeted therapies^47^. Heterogeneity of nucleic acid biomarkers is even higher for genetic diseases and autoimmune diseases. Rapid and affordable detection of the long tail of nucleic acid biomarkers is thus a general need for precision medicine for most if not all human disease.

The linear time complexity of DIMPLE enables high-plex primer design that scales to 10,000+ primers. This scalability enables redundant multiplex design (here, 5x degeneracy for each amplification target) that significantly reduced the experimental complexity of optimizing panels via primer replacement from predesigned pool. Design redundancy also allowed us to design primers without careful pre-selection of nonspecific genomic amplification, because primers contributing nonspecific genomic amplification can be replaced during the experimental optimization stage.

The multiplexing capability of a qPCR assay needs to be clarified as two separate concepts: *multiplicity of amplification* vs. *multiplicity of readout*. Beta thalassemia is a genetic disease caused by mutations in 3 genes (HBA1, HBA2, and HBB). Although over 300 pathogenic mutations have been documented in literature^48^, these can all be comprehensively covered by under 5 amplicons. Thus, beta thalassemia diagnostics is an application requiring low multiplicity of amplification and high multiplicity of readout. In contrast, for oncology gene fusion detection, such as the KMT2A and NTRK1/2/3 assays presented here, the actionability is a single binary decision: whether to use revumenib (for KMT2Ar+) or entrectinib (for NTRK1/2/3r+). Consequently, gene fusion detection for oncology therapy selection is an application that requires high multiplicity of amplification and low multiplicity of readout. Antimicrobial resistance detection applications falls somewhere in the middle, because different drug resistance plasmids and mutations contribute different degrees of resistance (quantitated as minimum inhibitory concentrations MIC^49^).

Combining DIMPLE primer design with high multiplicity of amplification and qPCR instrument platforms with high multiplicity of readout could unlock many molecular diagnostic applications heretofore only accessible via NGS, such as BRCA1/2 screening and homologous recombination deficiency (HRD) analysis for breast and ovarian cancer. Such assays could have significant advantages over NGS in rapid turnaround, lowered cost, ease-of-use, interpretability of data, and preservation of sensitive genetic information.

## MATERIALS AND METHODS

### DIMPLE Primer Design Strategy

Multiplex primer sets were designed using the DIMPLE algorithm. For each target, a context sequence flanking the region of interest (e.g., variant site or fusion breakpoint) was split into two Semiloci: the upstream (5′) sequence and the reverse complement of the downstream (3′) sequence. Primer candidates were generated by identifying subsequences meeting thermodynamic criteria: ΔG° of hybridization ≈ −12 kcal/mol (assuming 60°C annealing, 0.18 M Na⁺) and length of 15–35 nt. Primer selection maximized a composite FL score (Fitness − W × Loss). The Fitness score penalized homopolymers (>6 nt), suboptimal length, and self-complementarity. The Loss score estimated primer-dimer potential using k-mer hashing (6–10 nt), weighting reverse-complementary matches by length and GC content. A redundancy penalty deprioritized candidates for already-covered Semiloci until desired coverage (∼7 primers per target) was achieved.

### Oligonucleotides and Probes

PCR primers (IDT, Coralville, IA) were delivered at 100 µM in IDTE buffer (pH 8.0). TaqMan probes (IDT) were labeled with 5’ Cy5 and 3’ Iowa Black RQ quencher unless otherwise specified. All sequences are provided in **Supplementary Data 1**, organized by figure and assay type.

### Nucleic Acid Extraction and Preparation

Genomic DNA was extracted from pancreatic tissue (Prodo Labs, Aliso Viejo, CA) using the Monarch Genomic DNA Purification Kit (New England Biolabs, Ipswich, MA) and sheared to ∼250 bp using a Covaris M220 (Covaris, Woburn, MA). Total RNA was extracted from whole blood samples and cell lines using the Quick-RNA Whole Blood kit (Zymo Research, Irvine, CA). Concentration and quality were assessed by Qubit HS assays (Thermo Fisher Scientific, Waltham, MA). RNA was stored at −80°C; gDNA at −20°C, with ≤3 freeze-thaw cycles. For methylation assays, the Human Methylated & Non-Methylated (WGA) DNA Set (Zymo Research) served as reference material. Bisulfite conversion was performed using the EZ DNA Methylation-Lightning Kit (Zymo Research) with 1–10 ng input DNA.

### Gene Fusion Assay Workflow

For fusion detection assays, cDNA served as the qPCR template. cDNA was synthesized from up to 1000 ng total RNA using LunaScript RT SuperMix (New England Biolabs): 25°Cx2 min, 55°Cx10 min, 95°Cx1 min. Products were purified with 1.8× Ampure XP beads (Beckman Coulter, Brea, CA). For analytical validation, synthetic gBlocks (500 bp; IDT- see *Supplementary Data 1*) representing KMT2A fusion breakpoints were spiked into healthy donor cDNA at concentrations down to 0.05% VAF. Target copy number was verified by singleplex qPCR to within 2-fold of expected values – see *Supplementary Data 1*

### DIMPLE qPCR Assays

Reactions were performed on a CFX96 Touch (Bio-Rad) using PowerUp SYBR Green Master Mix (Thermo Fisher Scientific), except for methylation assays, which were performed on a QuantStudio 5 (Thermo Fisher Scientific) using UDG-free PrimeTime Gene Expression Master Mix (IDT) to preserve bisulfite-converted templates. Final primer concentrations were typically 15 nM; for KMT2A fusion assays, forward primers were used at 45 nM and reverse primers at 15 nM. Probes were used at 75–100 nM. Cycling: 95°C/5 min; 50 cycles of 95°C/30 s, 60°C/2 min. Analysis was performed using instrument software.

### NGS Library Preparation

To convert dUTP-containing amplicons to sequencing-compatible templates, DIMPLE products were purified (2.0× SPRI), then subjected to a 2-cycle rescue PCR using iTaq polymerase (Bio-Rad) with 5-min annealing at 60°C. Libraries were prepared using NEBNext Ultra II DNA Library Prep Kit (New England Biolabs), assessed by Bioanalyzer (Agilent, Santa Clara, CA) and Qubit, and sequenced (2 × 150 bp paired-end) on an Illumina MiSeq (Illumina, San Diego, CA).

### KMT2A Retrospective Studies

*US Cohort:* De-identified bone marrow samples (n=16) from patients with acute myeloid leukemia were obtained from MD Anderson Cancer Center (Houston, TX). As samples were de-identified retrospective remnants. *China Cohort:* Clinical samples (n=22) were collected and analyzed at Xinqiao Hospital (Chongqing, China); no biological material was transported internationally. All samples were de-identified prior to analysis.

### Data Analysis and Manuscript Drafting

Custom software for qPCR visualization, NGS primer dimer analysis, are available from the corresponding author upon request. Early drafts of this manuscript were generated with assistance from K-Dense AI by Biostate AI. All authors reviewed, edited, and take responsibility for the final content.

## Supporting information

DIMPLE_Supplemetary_Information

## DATA AVAILABILITY

The main data supporting the results in this study are available within the paper and the Supplementary Information. The raw and analyzed datasets generated during the study are available for research purposes from the corresponding author upon request.

## CODE AVAILABILITY

The custom software tools developed for this study—including the DIMPLE algorithm, the qPCR data visualization suite, and the NGS primer-dimer analysis pipeline—are proprietary and owned by Pupil Bio. All code is available from the corresponding author upon reasonable request for non-commercial research purposes.

## AUTHOR CONTRIBUTION

**Conceptualization:** G.C.I., D.Y.Z., A.P., X.Z. **Methodology:** G.C.I., D.Y.Z., A.P., X.Z. **Software:** A.P., D.Y.Z. **Investigation:** S.J.J., J.H., W.W., A.F., B.W., X.D., Q.W., C.Z. **Formal analysis:** G.C.I., D.Y.Z., X.Z., A.P., S.J.J., W.W., A.F., B.W., X.D., Q.W., C.Z. **Resources:** G.C.I., G.T., X.Z. **Writing – original draft:** G.C.I., A.P., D.Y.Z., X.Z. **Writing – review & editing:** All authors.

## ACKNOWLEDGEMENTS

We would like to express gratitude to Kellie Hull and Hannah Roberts for early experimental involvement.

## REFERENCES

1. Bustin, S. A. et al. The MIQE Guidelines: Minimum Information for Publication of Quantitative Real-Time PCR Experiments. Clin. Chem. 55, 611–622 (2009).

2. Schmitz, J. E., Stratton, C. W., Persing, D. H. & Tang, Y.-W. Forty Years of Molecular Diagnostics for Infectious Diseases. J. Clin. Microbiol. 60, e02446–21 (2022).

3. Khodakov, D., Wang, C. & Zhang, D. Y. Diagnostics based on nucleic acid sequence variant profiling: PCR, hybridization, and NGS approaches. Adv. Drug Deliv. Rev. 105, 3–19 (2016).

4. Goodwin, S., McPherson, J. D. & McCombie, W. R. Coming of age: ten years of next-generation sequencing technologies. Nat. Rev. Genet. 17, 333–351 (2016).

5. Boehme, C. C. et al. Rapid molecular detection of tuberculosis and rifampin resistance. N. Engl. J. Med. 363, 1005–1015 (2010).

6. Tansarli, G. S. & Chapin, K. C. Diagnostic test accuracy of the BioFire® FilmArray® meningitis/encephalitis panel: a systematic review and meta-analysis. Clin. Microbiol. Infect. 26, 281–290 (2020).

7. Khodakov, D., Li, J., Zhang, J. X. & Zhang, D. Y. Highly multiplexed rapid DNA detection with single-nucleotide specificity via convective PCR in a portable device. Nat. Biomed. Eng. 5, 702–712 (2021).

8. Sohaili, A. et al. Evaluating the effectiveness of FlashDx for diagnosing STIs among PrEP users in Kenya: A pilot study on diagnostic accuracy and prevalence. JMIR Prepr. 80643 (2025) doi:10.2196/preprints.80643.

9. Leber, A. L. et al. Multicenter Evaluation of the QIAstat-Dx Respiratory Panel for Detection of Viruses and Bacteria in Nasopharyngeal Swab Specimens. J. Clin. Microbiol. 58, e00155–20 (2020).

10. Elnifro, E. M., Ashshi, A. M., Cooper, R. J. & Klapper, P. E. Multiplex PCR: optimization and application in diagnostic virology. Clin. Microbiol. Rev. 13, 559–570 (2000).

11. Xie, N. G. et al. Designing highly multiplex PCR primer sets with Simulated Annealing Design using Dimer Likelihood Estimation (SADDLE). Nat. Commun. 13, 1881 (2022).

12. Rachlin, J., Ding, C., Cantor, C. & Kasif, S. Computational tradeoffs in multiplex PCR assay design for SNP genotyping. BMC Genomics 6, 102 (2005).

13. Mamanova, L. et al. Target-enrichment strategies for next-generation sequencing. Nat. Methods 7, 111–118 (2010).

14. Shen, Z. et al. MPprimer: a program for reliable multiplex PCR primer design. BMC Bioinformatics 11, 143 (2010).

15. Wingo, T. S., Kotlar, A. & Cutler, D. J. MPD: multiplex primer design for next-generation targeted sequencing. BMC Bioinformatics 18, 14 (2017).

16. Kechin, A., Borisova, U., Blinov, A. & Filipenko, M. NGS-PrimerPlex: high-throughput primer design for multiplex polymerase chain reactions. PLoS Comput. Biol. 16, e1008468 (2020).

17. Kirkpatrick, S., Gelatt, C. D. Jr. & Vecchi, M. P. Optimization by simulated annealing. Science 220, 671–680 (1983).

18. SantaLucia, J. Jr. & Hicks, D. The thermodynamics of DNA structural motifs. Annu. Rev. Biophys. Biomol. Struct. 33, 415–440 (2004).

19. Zhang, J. X. et al. Predicting DNA hybridization kinetics from sequence. Nat. Chem. 10, 91–98 (2018).

20. Untergasser, A. et al. Primer3—new capabilities and interfaces. Nucleic Acids Res. 40, e115 (2012).

21. AACR Project GENIE Consortium. AACR Project GENIE: Powering Precision Medicine through an International Consortium. Cancer Discov. 7, 818–831 (2017).

22. Issa, G. C. et al. Therapeutic implications of menin inhibition in acute leukemias. Leukemia 35, 2482–2495 (2021).

23. Meyer, C. et al. The KMT2A recombinome of acute leukemias in 2023. Leukemia 37, 988–1005 (2023).

24. Issa, G. C. et al. The menin inhibitor revumenib in KMT2A-rearranged or NPM1-mutant leukaemia. Nature 615, 920–924 (2023).

25. Wang, E. S. et al. Ziftomenib in relapsed or refractory acute myeloid leukaemia (KOMET-001): a multicentre, open-label, multi-cohort, phase 1 trial. Lancet Oncol. 25, 1310–1324 (2024).

26. Issa, G. C. et al. Menin Inhibition With Revumenib for KMT2A-Rearranged Relapsed or Refractory Acute Leukemia (AUGMENT-101). J. Clin. Oncol. 43, 75–84 (2025).

27. Issa, G. C. et al. Predictors of outcomes in adults with acute myeloid leukemia and KMT2A rearrangements. Blood Cancer J. 11, 162 (2021).

28. Zeidner, J. F. et al. Azacitidine, Venetoclax, and Revumenib for Newly Diagnosed NPM1-Mutated or KMT2A-Rearranged AML. J. Clin. Oncol. 43, 2606–2615 (2025).

29. Döhner, H. et al. Diagnosis and management of AML in adults: 2017 ELN recommendations from an international expert panel. Blood 129, 424–447 (2017).

30. Issa, G. C. et al. A Novel NGS Assay to Detect Any KMT2A fusion Transcript at Low Levels. Blood 140, 10738–10740 (2022).

31. Heuser, M. et al. 2021 Update on MRD in acute myeloid leukemia: a consensus document from the European LeukemiaNet MRD Working Party. Blood 138, 2753–2767 (2021).

32. Kim, P. & Zhou, X. FusionGDB: fusion gene annotation DataBase. Nucleic Acids Res. 47, D994–D1004 (2019).

33. Doebele, R. C. et al. Entrectinib in patients with advanced or metastatic NTRK fusion-positive solid tumours: integrated analysis of three phase 1-2 trials. Lancet Oncol. 21, 271–282 (2020).

34. Drilon, A. et al. Efficacy of Larotrectinib in TRK Fusion-Positive Cancers in Adults and Children. N. Engl. J. Med. 378, 731–739 (2018).

35. Westphalen, C. B. et al. Genomic context of NTRK1/2/3 fusion-positive tumours from a large real-world population. Npj Precis. Oncol. 5, 69 (2021).

36. Thway, K. & Fisher, C. Mesenchymal Tumors with EWSR1 Gene Rearrangements. Surg. Pathol. Clin. 12, 165–190 (2019).

37. Tsuda, Y. et al. The clinical heterogeneity of round cell sarcomas with EWSR1/FUS gene fusions: Impact of gene fusion type on clinical features and outcome. Genes. Chromosomes Cancer 59, 525–534 (2020).

38. Hensel, T. et al. BET bromodomain inhibitors suppress EWS-FLI1-dependent transcription and the IGF1 autocrine mechanism in Ewing sarcoma. Oncotarget 7, 43504–43517 (2016).

39. Sankar, S. et al. Reversible LSD1 inhibition interferes with global EWS/ETS transcriptional activity and impedes Ewing sarcoma tumor growth. Clin. Cancer Res. 20, 4584–4597 (2014).

40. Jones, P. A. Functions of DNA methylation: islands, start sites, gene bodies and beyond. Nat. Rev. Genet. 13, 484–492 (2012).

41. Bird, A. DNA methylation patterns and epigenetic memory. Genes Dev. 16, 6–21 (2002).

42. Moss, J. et al. Comprehensive human cell-type methylation atlas reveals origins of circulating cell-free DNA in health and disease. Nat. Commun. 9, 5068 (2018).

43. Loyfer, N. et al. A DNA methylation atlas of normal human cell types. Nature 613, 355–364 (2023).

44. Frommer, M. et al. A genomic sequencing protocol that yields a positive display of 5-methylcytosine residues in individual DNA strands. Proc. Natl. Acad. Sci. 89, 1827–1831 (1992).

45. Vaisvila, R. et al. Enzymatic methyl-seq detects DNA methylation at single-base resolution from picograms of DNA. Genome Res. 31, 1280–1289 (2021).

46. Larsson, D. G. J. & Flach, C.-F. Antibiotic resistance in the environment. Nat. Rev. Microbiol. 20, 257–269 (2022).

47. Dagogo-Jack, I. & Shaw, A. T. Tumour heterogeneity and resistance to cancer therapies. Nat. Rev. Clin. Oncol. 15, 81–94 (2018).

48. Giardine, B. M. et al. Clinically relevant updates of the HbVar database of human hemoglobin variants and thalassemia mutations. Nucleic Acids Res. 49, D1192–D1196 (2021).

49. Kowalska-Krochmal, B. & Dudek-Wicher, R. The Minimum Inhibitory Concentration of Antibiotics: Methods, Interpretation, Clinical Relevance. Pathogens 10, 165 (2021).

