## Supplementary material for "Scaling Multiplex qPCR Primer Design to 1000-plex using the Degenerate Incomplete Multiplex Primer List Extension (DIMPLE) Algorithm": DIMPLE_Supplemetary_Information

**Section S1 | List of primers used**

All primer sequences, probes, and synthetic spike-ins used in this work are provided in Supplementary Data 1, organized by assay in dedicated tabs corresponding to main text figures: 100plex DIMPLE (*Figure 1b*), 100plex Primer3 (*Figure 1c*), 1000plex Assay (*Figure 2*), KMT2A Assay (*Figures 3–4*), NTRK Assay (*Figure 5a*), EWSR1_FUS Assay (*Figure 5b*), and Methylation Panel (*Figure 6*). Each tab contains five columns: sequence name, type (FW, RV, TQ, or gBlock), sequence, gene, and target BED coordinate or exon.

**Section S2 | 1000-plex DIMPLE design**

***Loci Selection.*** The panel was designed to amplify 1,000 loci encompassing a total of 22,712 oncogenic variants across 364 genes identified from the AACR Project GENIE® database^[1]^ and associated with colorectal, breast, ovarian, lung, and pancreatic (both PDAC and non-PDAC) tumors. A summary of the panel is provided in *Supplementary Data 2*, containing four columns: gene symbol, BED coordinates (GRCh38), context sequence (BED ± 150 nt, with mutation positions in uppercase), and list of covered mutations.

The 1,000 loci were selected to maximize the number of covered cases in non-overlapping regions, with each region constrained to ≤70 bp in length to ensure sufficient design space for amplicons compatible with a variety of sample types, including cell-free DNA (cfDNA), which typically requires amplicons shorter than ~169 nt.

***Primer Design.*** Primers were designed using the DIMPLE software with a maximum amplicon length of 170 bp and a redundancy of 7 (allowing up to seven interchangeable primer versions per locus). Input parameters included an acceptable amplicon length range of 70–170 bp, an annealing temperature of 60°C, and salt concentration adjusted for the PCR master mix used. No minimum primer length was enforced; maximum primer length was set to 40 nt. For the 1000-plex proof-of-concept, primers were not screened by BLAST, as the goal was to maximize insight into primer performance relative to dimer formation. All 11,627 designed sequences are reported in *Supplementary Data 1*, tab *1000plex Assay*. Primers used in each experiment are indicated by TRUE values in the corresponding figure columns (Fig2B, Fig2D, Fig2E).

A subset of 200 primers (100 targets; *Supplementary Data 1*, tab *100plex DIMPLE*) was randomly selected and pooled in a 100-plex assay to benchmark against Primer3-designed counterparts (*Supplementary Data 1*, tab 100plex *Primer3*).

. Primer3 was run via primer3-py (<https://github.com/libnano/primer3-py> ); as the software lacks native multiplex capability, primers for each target were designed independently without cross-target interaction screening. The top-ranked primer pair for each target was selected and ordered. Design parameters were matched to those used for DIMPLE.

**Section S3 | NGS Data Analysis**

To evaluate the 1,000-plex assay, two libraries were prepared: a no-template control (**NTC**) and a positive control (**PC**), sequenced at 0.2M and ~4M read depth, respectively. The NTC reaction was used directly in final indexing PCR (9 cycles), while PC was diluted 1:100 before indexing (6 cycles). The lower **NTC** depth prevents cluster poisoning; the higher **PC** depth enables assessment of on-target coverage distribution.

Raw FASTQ files were processed using VISIO (Verified Integrated Sequencing and Interpretative Outcomes), a custom bioinformatics pipeline integrating Bowtie2 alignment with Python scripts for alignment validation and primer-dimer identification. The pipeline first trims adapter sequences (Cutadapt), aligns paired-end reads to the reference genome (Bowtie2), and processes BAM files (samtools). Reads are retained if they meet the following criteria: alignment length ≥80 bp, ≤2 mismatches, indel length ≤2 bp, concordant mate-pairs, and ≥95% amplicon coverage. Given that all amplicons in this study are <150 bp, these thresholds effectively distinguish on-target amplification from artifacts.

Reads failing any criterion are routed to the dimer analysis module, which collapses identical sequences and identifies primer sequences using Levenshtein distance matching (≤4 mismatches allowed). Artifacts are then classified as dimers (two primer sequences with ≥1 bp overlap), byproducts (two primers identified but non-overlapping, typically representing off-target amplification or concatemers), or unidentified (one or both primers not detected, typically NGS library artifacts). Finally, the pipeline simulates iterative primer-pair elimination to rank candidates for removal or substitution based on their contribution to total dimer signal.

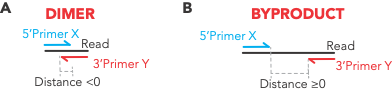

**Supplementary Figure 1 Classification of primer artifacts**. (A) Dimers: forward and reverse primers overlap (distance <0). (B) Byproducts: primers are non-overlapping (distance ≥0), indicating intervening sequence from off-target amplification or concatemerization.

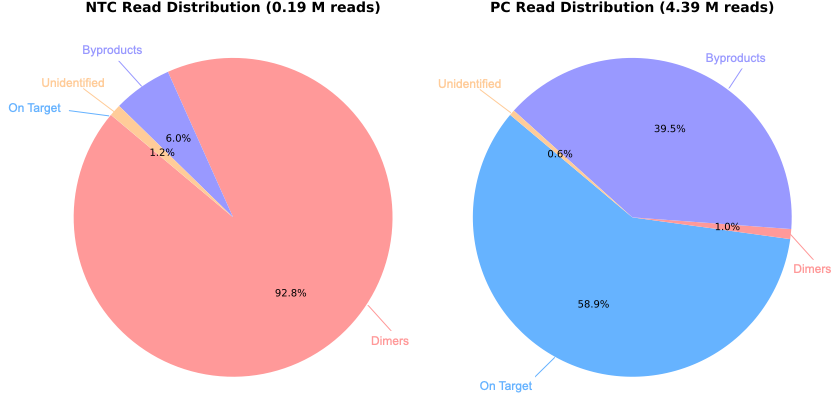

**Supplementary Figure 2 Read distribution in the 1,000-plex DIMPLE assay.** Classification of sequencing reads from no-template control (NTC) and positive control (PC) libraries. NTC is dominated by dimers (92.8%); PC shows 58.9% on-target reads with byproducts (39.5%) as the primary artifact.

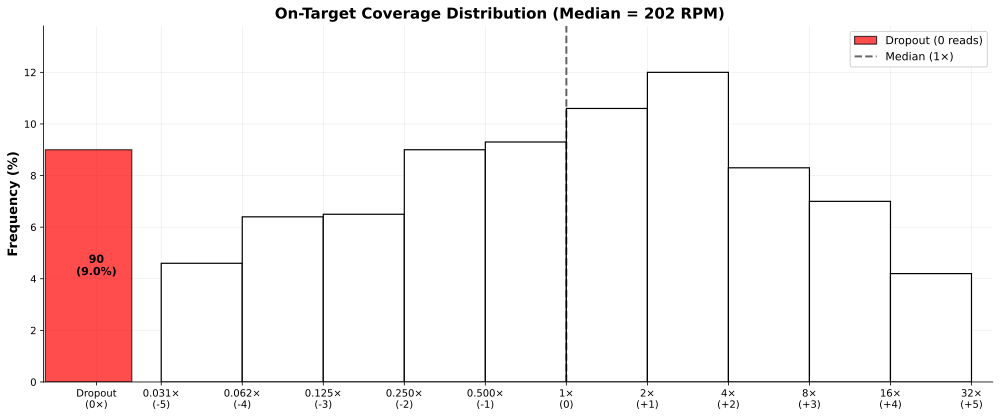

**Supplementary Figure 3 On-target coverage distribution for the Positive Control sample**. Histogram showing amplicon coverage normalized to median (202 RPM). Red bar indicates dropout amplicons (0 reads; 9.0%). Dashed line marks median coverage (1×). X-axis shows fold-change relative to median on a log₂ scale.

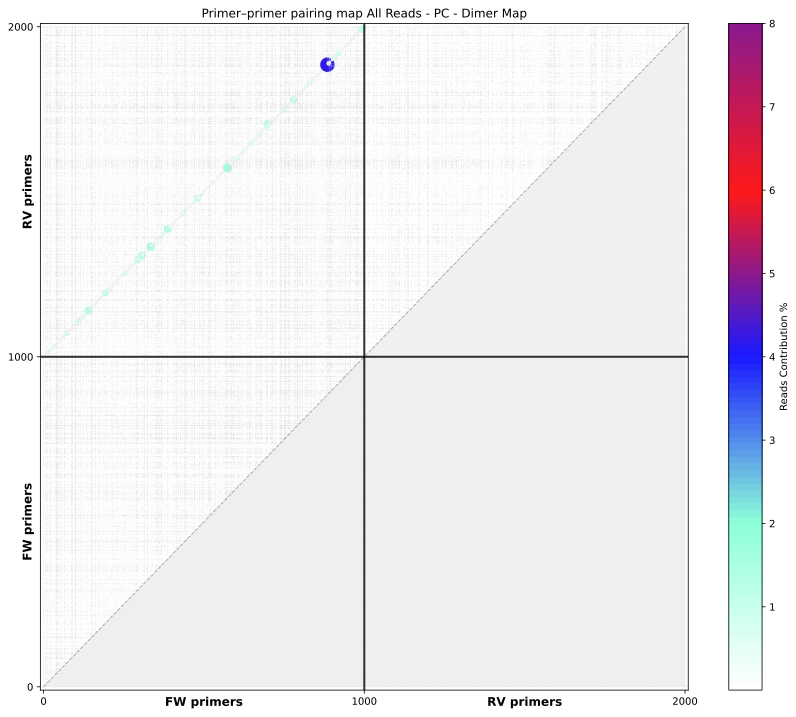

**Supplementary Figure 4 Primer–primer pairing map for the positive control – all reads.** Heatmap showing pairwise primer interactions, with color intensity indicating read contribution (%). Points along the diagonal represent correctly paired forward and reverse primers targeting the same amplicon. The majority of reads cluster on the diagonal, confirming that most sequenced products arise from intended primer pairs.

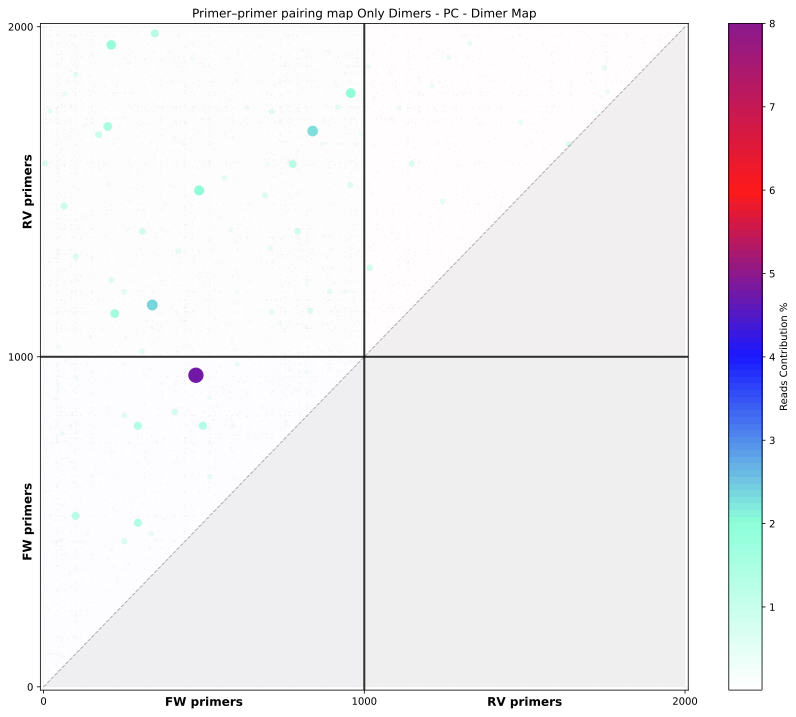

**Supplementary Figure 5 Primer–primer pairing map for dimer-forming reads in the positive control.** Heatmap showing only reads classified as dimers, with color intensity indicating read contribution (%). Off-diagonal clustering identifies specific primer pairs prone to dimer formation, guiding targeted primer optimization.

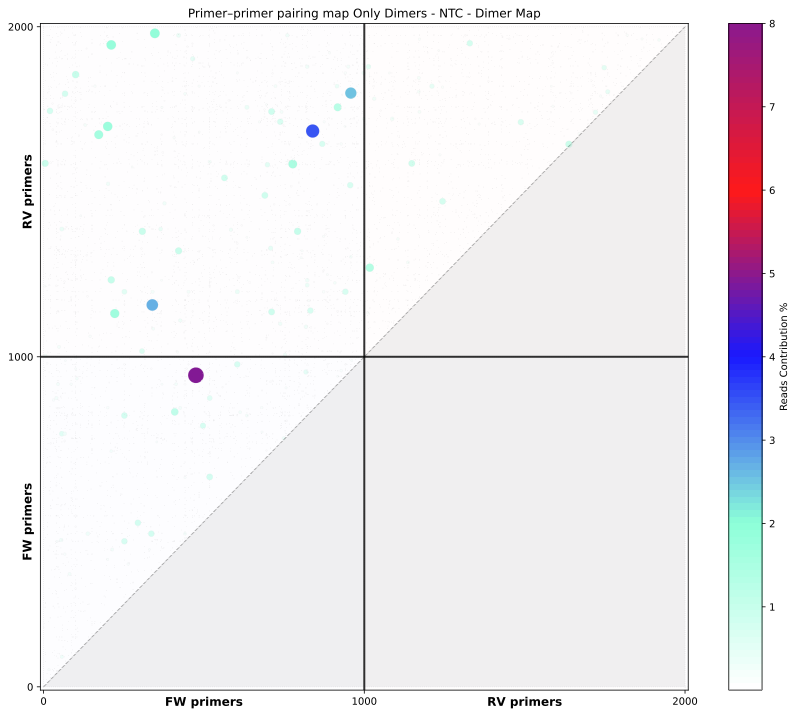

**Supplementary Figure 6 Primer–primer pairing map for dimer-forming reads in the NTC.** Heatmap showing only reads classified as dimers, with color intensity indicating read contribution (%). Off-diagonal clustering identifies specific primer pairs prone to dimer formation, guiding targeted primer optimization.

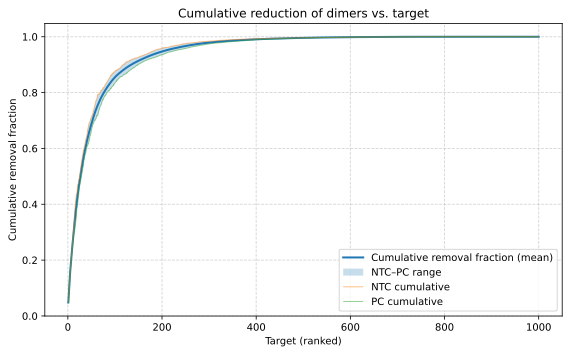

**Supplementary Figure 7 Cumulative dimer reduction by iterative primer pair elimination.** Simulated removal of primer pairs ranked by their contribution to total dimer reads. Curves show cumulative fraction of dimers eliminated for NTC (orange), PC (green), and mean (blue); shaded area indicates NTC–PC range. 38 Primers pairs account for ~60% of the dimers, while the next 124 primer pairs account for an additional 30% of dimers.

**Section S4 | KMT2 Assay Design**

We designed a multiplex panel targeting 12 *KMT2A* fusion partners (*AFDN, AFF1, ELL, EPS15, MLLT1, MLLT10, MLLT11, MLLT3, MLLT6, SEPTIN6, SEPTIN9,* and *USP2*), covering approximately 90% of *KMT2A*-rearranged leukemias and >95% of actionable fusions (excluding partial tandem duplications). Focusing on the Major and Minor *KMT2A* breakpoints where the 5' region is retained, we utilized the DIMPLE algorithm to design 12 forward primers targeting critical *KMT2A* exons and 192 reverse primers targeting the canonical exons of the fusion partners (excluding UTRs), as detailed in *Supplementary Table 1*.

The design employed a 2× redundancy strategy. To ensure robust detection and accommodate TaqMan probes, forward primers were positioned >20 nt from the junction, reverse primers >5–10 nt from exon boundaries, and amplicon lengths were constrained to <150 bp to maximize sensitivity; all other parameters were maintained as described above.

| Gene | Role | Number of Exons Targeted |
| --- | --- | --- |
| KMT2A | Target (5' Anchor) | 12 |
| AFDN | Partner | 33 |
| AFF1 | Partner | 21 |
| ELL | Partner | 12 |
| EPS15 | Partner | 25 |
| MLLT1 | Partner | 11 |
| MLLT10 | Partner | 22 |
| MLLT11 | Partner | 2 |
| MLLT3 | Partner | 11 |
| MLLT6 | Partner | 19 |
| SEPTIN6 | Partner | 11 |
| SEPTIN9 | Partner | 12 |
| USP2 | Partner | 13 |
| Total |  | 204 |

**Supplementary Table 1 Genes and number of Exons included in the KMT2A panel.**

**Section S5 | KMT2 Assay Analytical Validation**

**Synthetic Spike-ins:** Background matrix was prepared from whole blood collected by a certified phlebotomist and processed within 24 hours. RNA was extracted and reverse-transcribed as described in *Materials and Methods*.

Wild-type KMT2A cDNA was quantified using primers targeting exon 2 (FW: CAATTCTTAGGTTTGGCTCAGATG; RV: AAGGAGACCTTGTGGGACTTC), and calibrated against pre-characterized human genomic DNA. Synthetic gBlocks were spiked into a background of 10,000 wild-type KMT2A copies and verified to be within 2-fold of expected concentrations. Individual titration curves are shown in *Supplementary Figure 8*.

**Cell Line Validation**. To de-risk assay performance on clinical samples, we performed an independent validation using biological material. RNA was extracted from the KMT2Ar(+) cell line RS4;11 (CRL-1873; ATCC) and from KMT2Ar(−) leukocytes (Takara Bio) using the Monarch Total RNA Miniprep Kit (New England Biolabs). Positive RNA was then serially diluted into the wild-type RNA matrix to generate samples representing VAFs from 10% down to 0.001%. Unlike synthetic spike-ins, this approach captures the full complexity of biological samples, including potential matrix effects from competing transcripts. Samples were reverse-transcribed and analyzed as described in Materials and Methods. The assay detected KMT2A::AFF1 down to 0.01% VAF, consistent with synthetic validation results (*Supplementary Figure 9*).

| 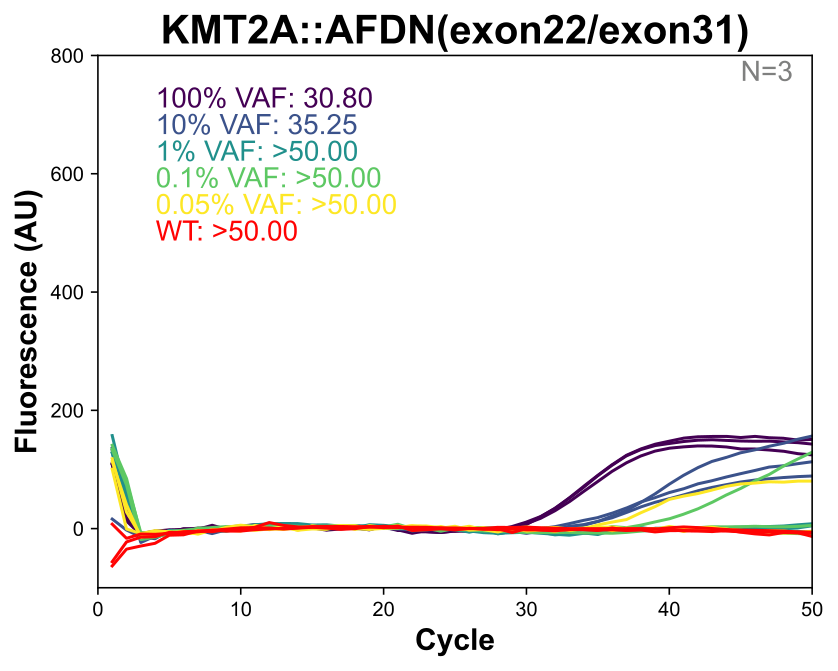 | 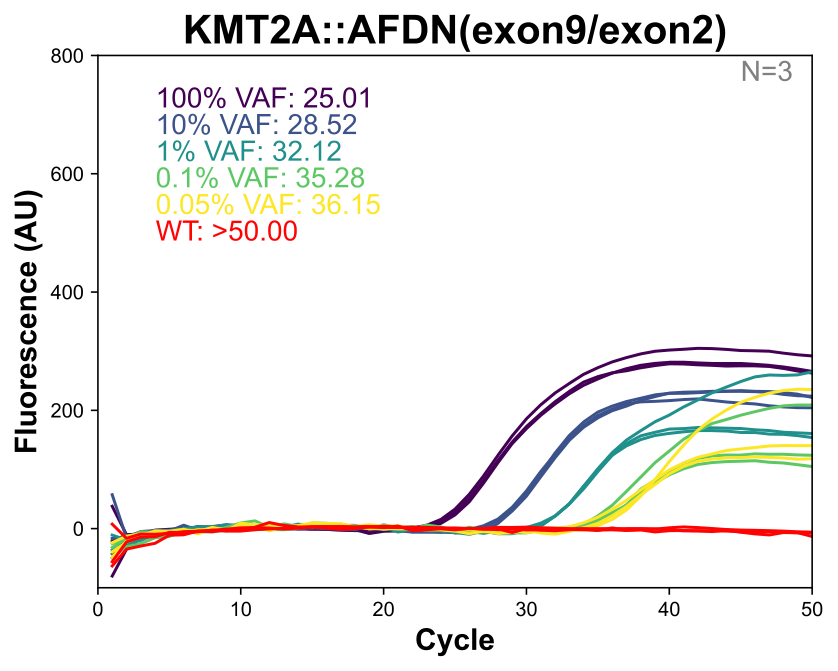 |
| --- | --- |
| 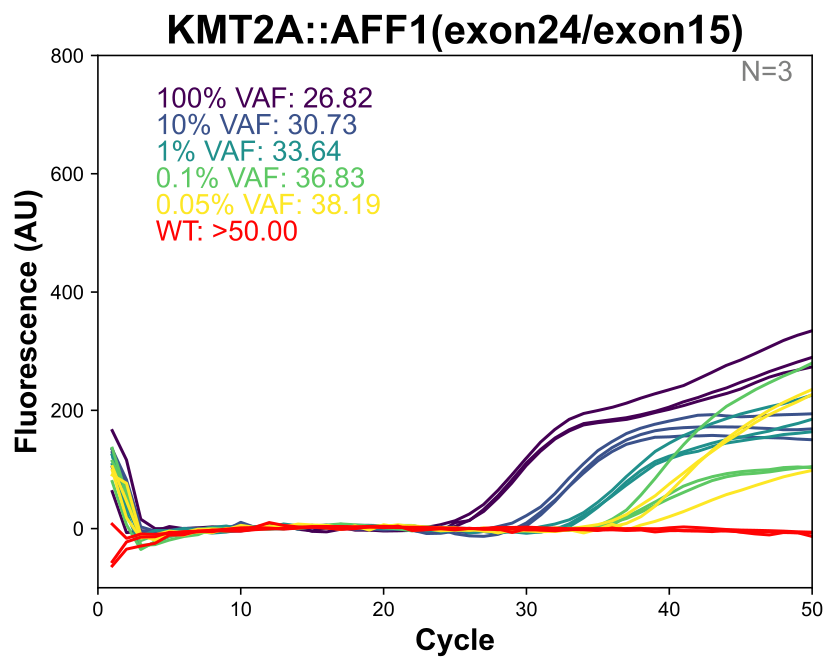 | 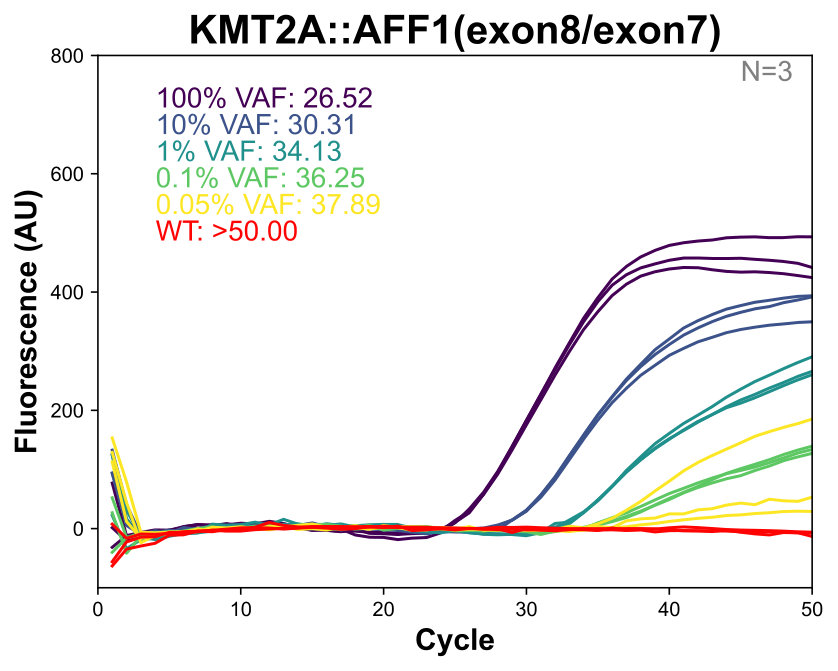 |
| 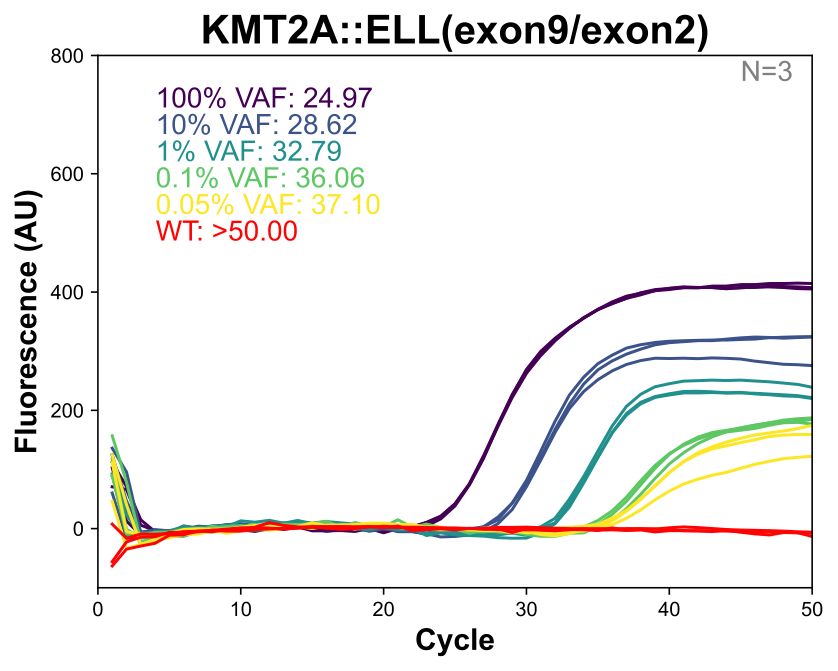 | 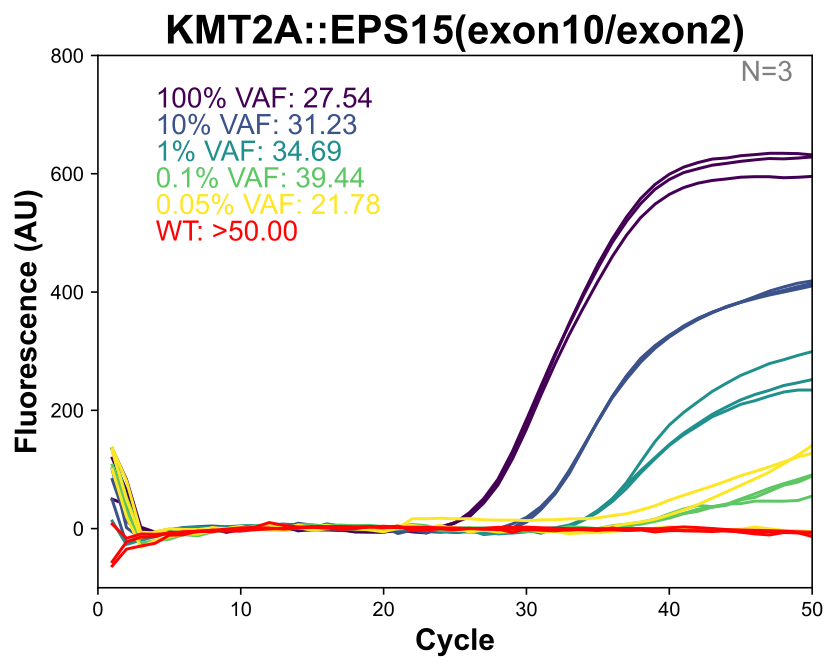 |

| 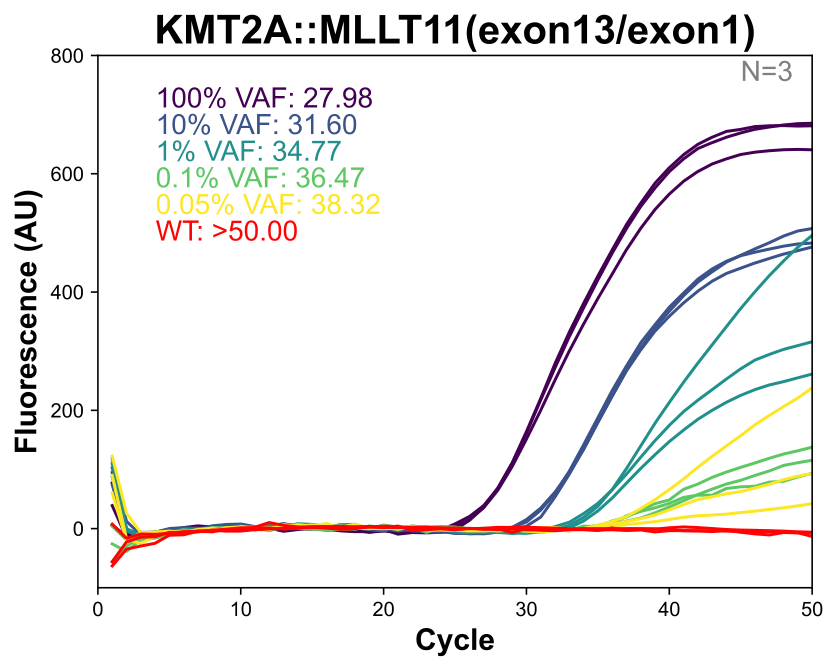 | 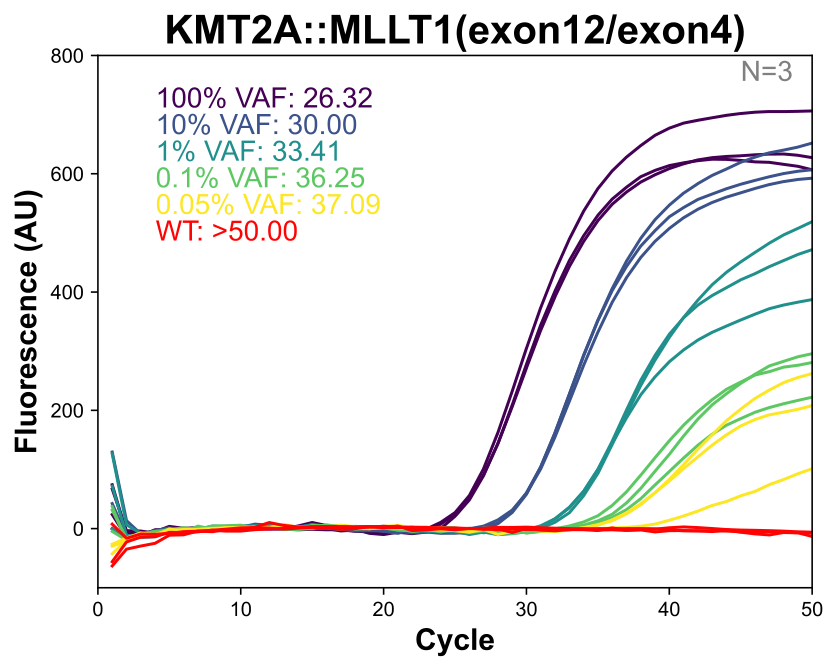 |
| --- | --- |
| 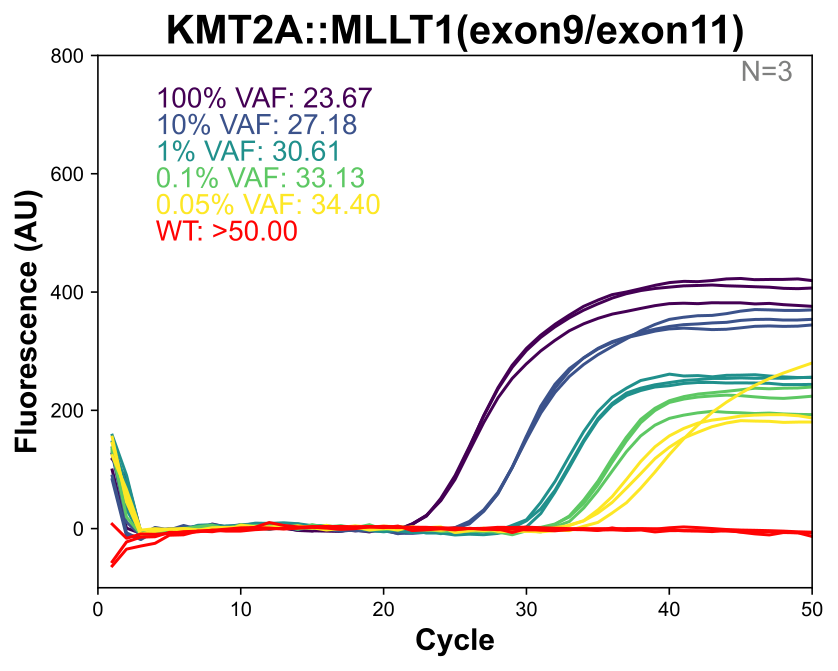 | 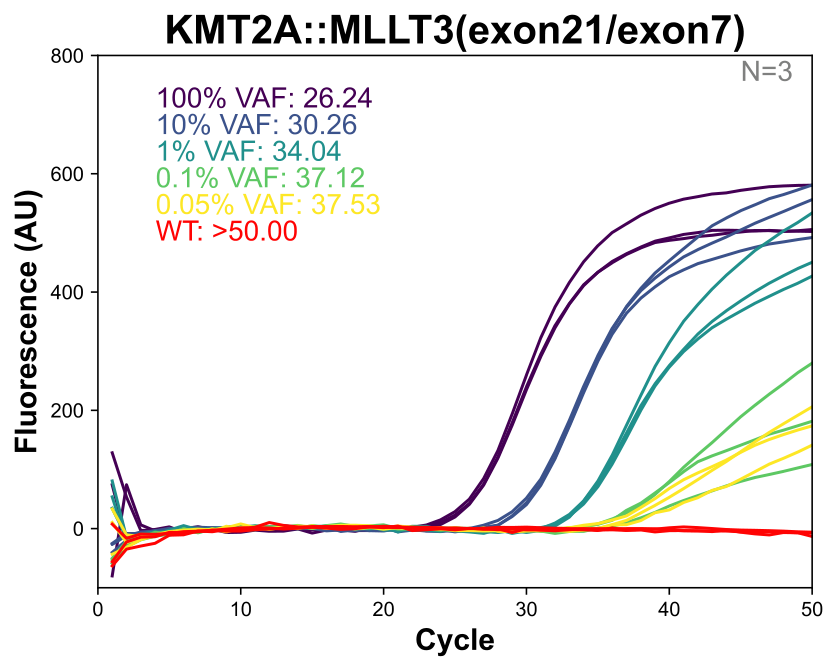 |
| 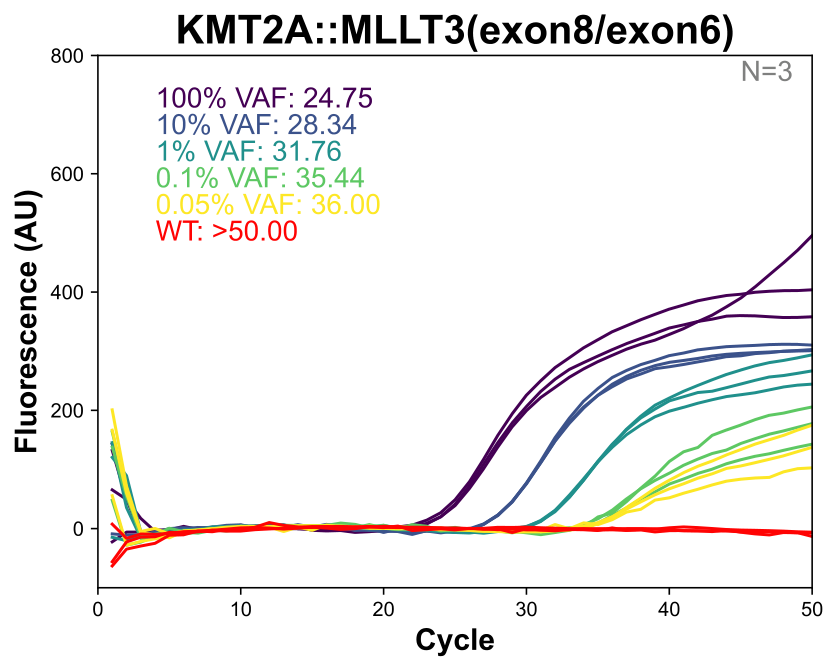 | 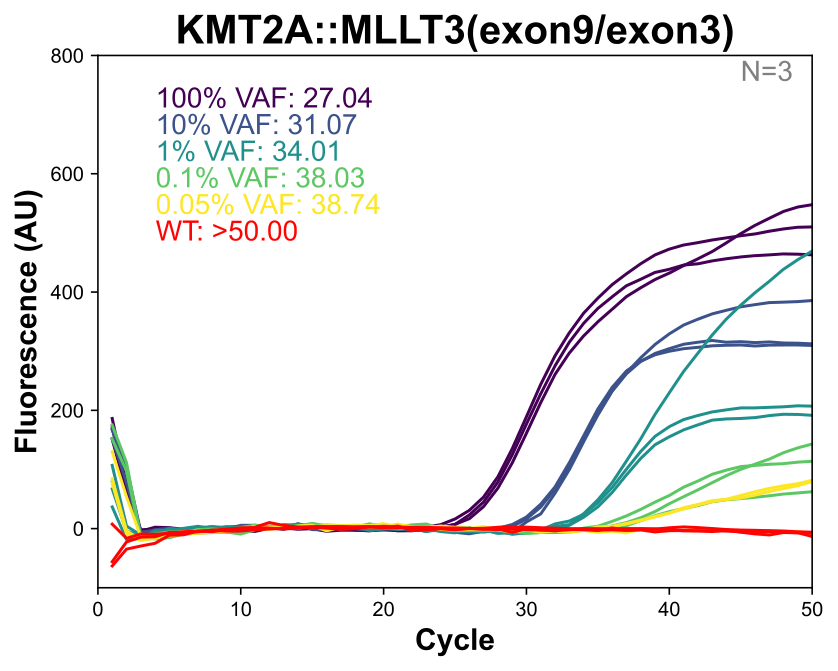 |

| 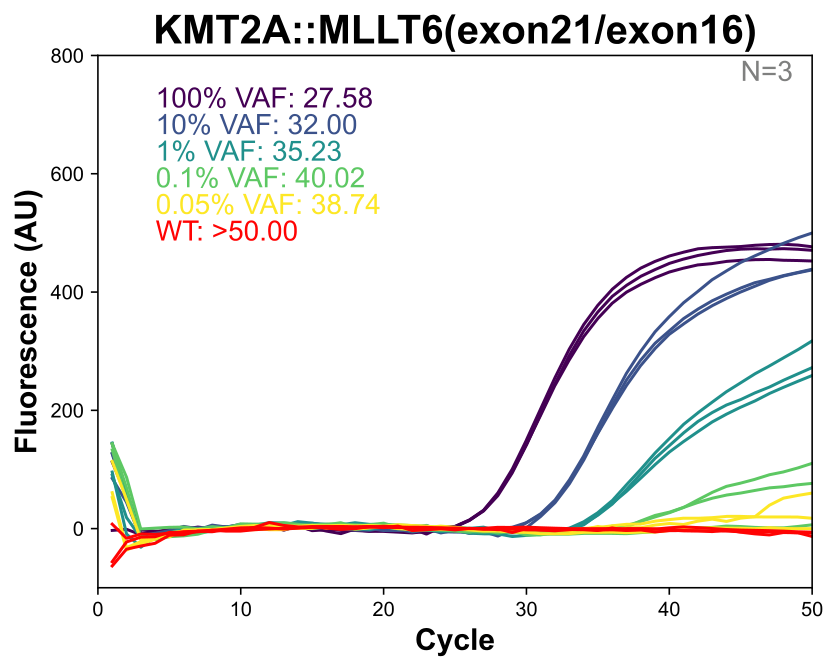 | 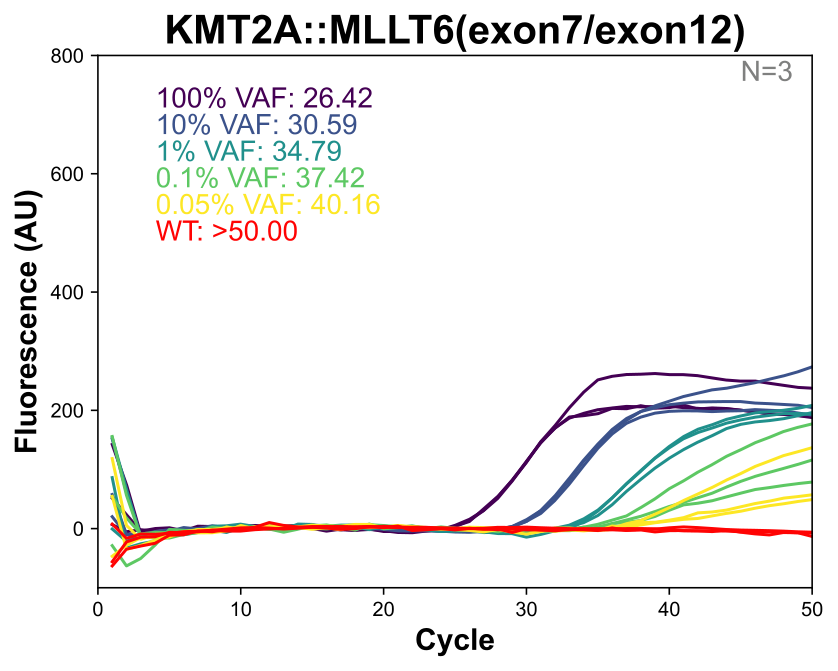 |
| --- | --- |
| 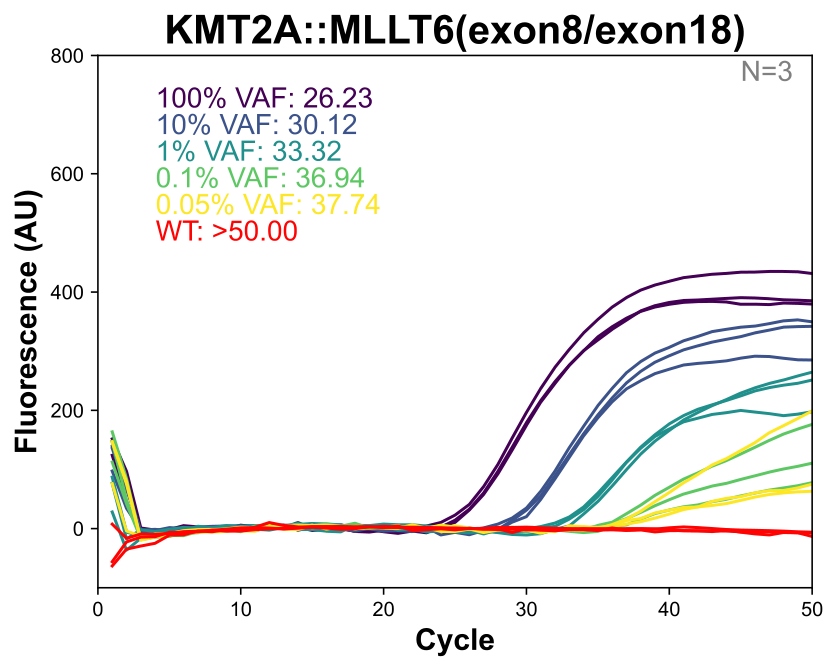 | 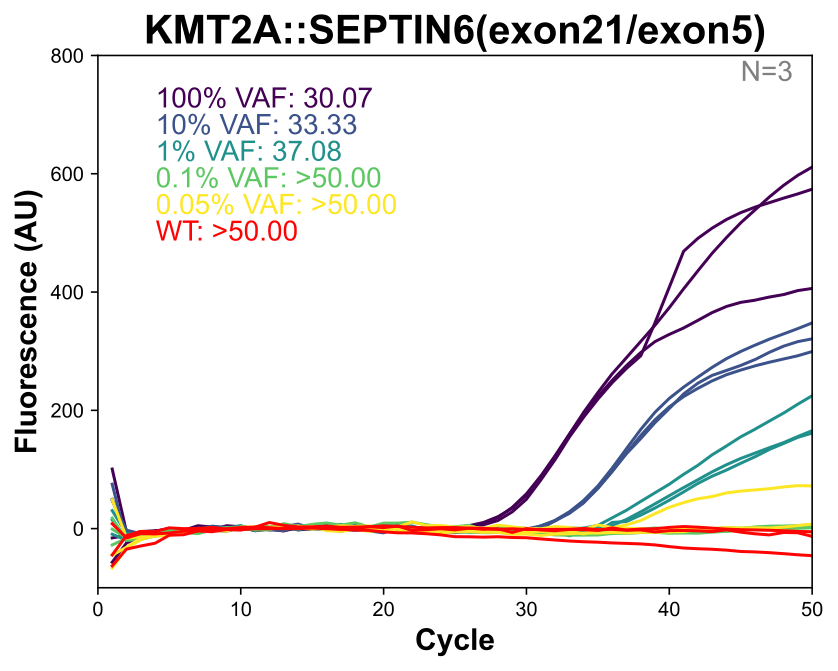 |
| 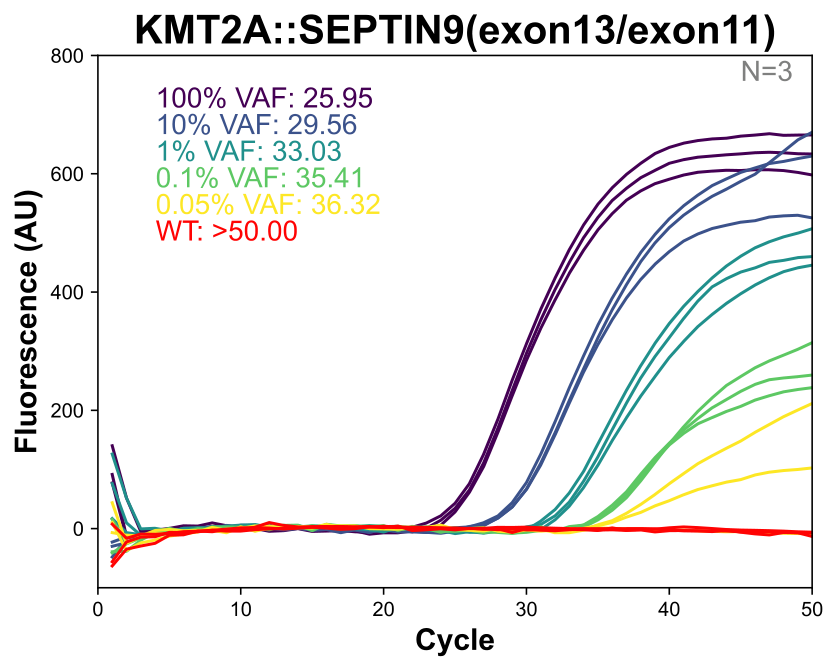 | 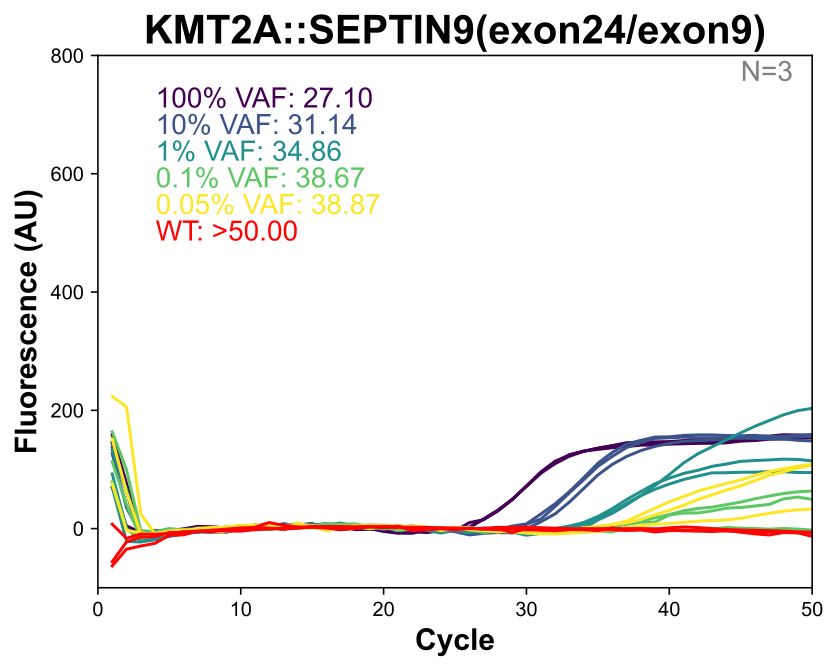 |

| 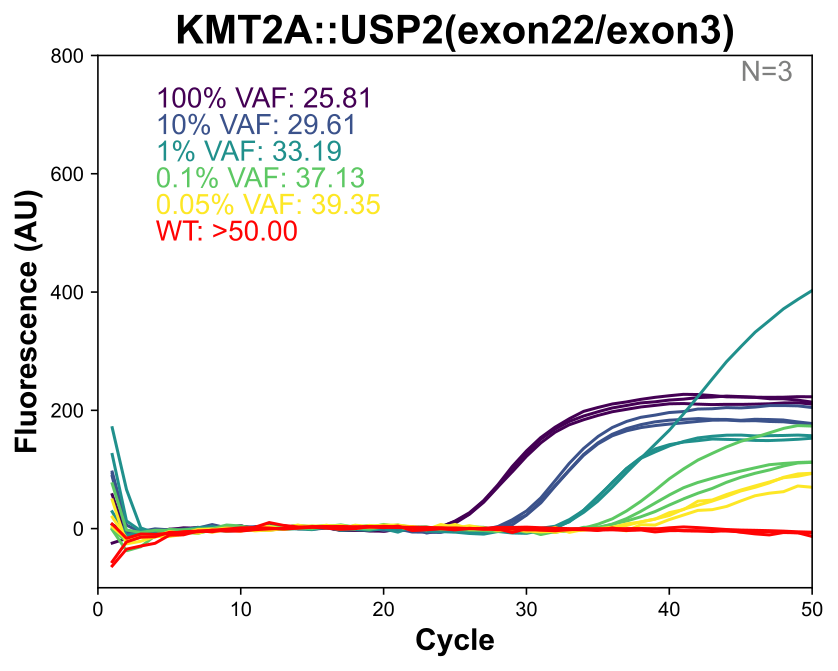 |
| --- |

**Supplementary Figure 8 qPCR titration curves for 19 KMT2A fusion targets**. Each panel shows amplification curves for a single fusion across serial dilutions from 100% to 0.05% VAF. Panels are labeled by fusion partner gene.

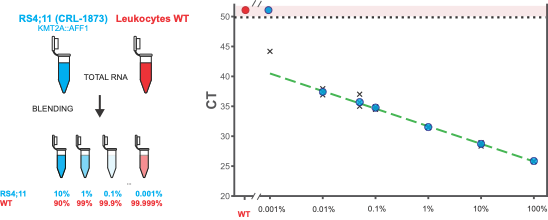

**Supplementary Figure 9 KMT2A::AFF1 fusion detection in blended RNA samples.** (Left) Blending schematic. (Right) Cq values across VAF dilutions. Blue circles: fusion signal; black X: replicates; dashed line: linear fit. Shaded region: no amplification (Cq ≥50).

**Section S6 | MD Anderson Cohort Characteristics and Assay Results**

**Cohorts and Demographics** We analyzed a retrospective cohort of 16 blinded AML samples collected at MD Anderson Cancer Center. The baseline demographic and clinical characteristics of the patients are summarized in *Supplementary Table 2.*

| Characteristic | Value |
| --- | --- |
| Median age (years) | 62 |
| Age range | 43-76 |
| Female | 8 |
| Female % | 50% |
| Blast % median | 56 |
| Blast % range | 12-92 |
| AML | 12 |
| AML % | 75% |
| AMOL | 4 |
| AMOL % | 25% |
| Medullary and extramedullary | 0 |
| Abnormal 11q | 9 |
| Abnormal 11q % | 56% |
| Trisomy 8 | 1 |
| Trisomy 8 % | 6% |
| -5/5q- | 4 |
| -5/5q- % | 25% |
| Diploid | 1 |
| Diploid % | 6% |

**Supplementary Table 2 Baseline characteristics of patients whose samples were selected for the study.**

**Cytogenetic and FISH Classifications.** To evaluate assay performance in the context of complex genomic landscapes, we examined the full clinical cytogenetic and FISH reports for all samples (*Supplementary Table 3*). Several samples exhibited complex karyotypes where cytogenetics and FISH provided complementary information. Sample **MD16** displayed monosomy 11 (−11) by conventional karyotyping, while FISH reported KMT2A amplification localized to marker chromosomes—both consistent with chromosomal instability without evidence of a targetable fusion. Sample **MD7** was characterized by cytogenetics as harboring a t(4;11) translocation, while the clinical FISH report described a "3′ deletion" of the KMT2A signal; these findings are not mutually exclusive, as complex rearrangements can produce atypical FISH patterns.

In contrast, sample **MD15** represented a true methodological discrepancy: cytogenetics revealed a t(2;11) translocation (KMT2A::THADA) but tested negative by standard break-apart FISH, likely reflecting the inherent limitations of FISH probe design for rare fusion partners. Notably, our DIMPLE assay also reported **MD15** as negative, as THADA is not among the 12 partner genes included in the current panel design.

| **ID** | **Cytogenetics** | **FISH** | **BM Blast %** |
| --- | --- | --- | --- |
| MD1 | 42,XX,dup(1)(q23q32),-3,der(5;17)(p10;q10),-7,add(9)(q22),-20[6]/42,idem,+mar[cp8]/84,idemx2,+1~3mar[cp4]/46,XX,inv(3)(p21q12)[1]/46,XX[1] | nuc ish(KMT2Ax4)[16/200] | 72 |
| MD2 | 46,XY,t(11;17)(q23;q25)[20] | nuc ish(KMT2Ax2)(5'KMT2A sep 3'KMT2Ax1)[166/200] | 41 |
| MD3 | 43,XX,der(1;5)(q10;p10),add(2)(q37),+8,add(12)(p11.2),-17,-18,-20[2]/44,idem,+mar[14]/47,idem,add(3)(p21),+17,+18,+20,+mar[1]/81~87,idemx2,+2~4mar[cp3] | nuc ish(KMT2Ax4)[13/200] | 72 |
| MD4 | 48,XX,+8,+11[20] | nuc ish(KMT2Ax3)[186/200] | 41 |
| MD5 | 46,XX,t(9;11)(p22;q23)[?] | ND | 77 |
| MD6 | 47,XX,+8,t(9;11)(p22;q23)[10] | ND | 81 |
| MD7 | 84~87,XXYY,-1,der(4)t(4;11)(q21;q23),t(4;11),-5,-10,-11,-21[cp20] | nuc ish(5'KMT2Ax3,3'KMT2Ax2)(5'KMT2A con 3'KMT2Ax2)[181/200]/(5'KMT2Ax2,3'KMT2Ax1)(5'KMT2A con 3'KMT2Ax1)[10/200] | 85 |
| MD8 | 47,XY,inv(9)(p12q13),t(9;11)(p22;q23),+21[12]//46,XX[8] | nuc ish(KMT2Ax2)(5'KMT2A sep 3'KMT2Ax1)[184/200] | 80 |
| MD9 | 42~45,XX,add(5)(q11.2),-9,-9,add(15)(p13),add(17)(p11.2),der(17)add(17)(p13)dup(17)(p13q12),-19,-21,-21,-22,+2~6mar[cp17]/46,XX[3] | nuc ish(KMT2Ax4)[13/200]/(KMT2Ax3)[12/200] | 21 |
| MD10 | 42~44,X,-Y,dic(1;17)(p32;p13),add(4)(p12),add(8)(q24.3),add(11)(p11.2),-13,add(13)(q34),-16,t(16;21)(q24;q22),add(18)(q23),del(19)(p13.3),-21,+1~2mar[cp20] | nuc ish(KMT2Ax3)[30/200] | 68 |
| MD11 | 46,XY,t(11;19)(q23;p13.1)[19]/46,XY[1] | nuc ish(KMT2Ax2)(5'KMT2A sep 3'KMT2Ax1)[183/200] | 28 |
| MD12 | 46,XY,t(11;19)(q23;p13.3)[20] | nuc ish(KMT2Ax2)(5'KMT2A sep 3'KMT2Ax1)[160/200] | 89 |
| MD13 | 46,XY[20] | nuc ish(KMT2Ax2)[200] | 14 |
| MD14 | 46,XX,t(9;11)(p22;q23)[20] | nuc ish(KMT2Ax2)(5'KMT2A sep 3'KMT2Ax1)[184/200] | 51 |
| MD15 | 46,XX,t(2;11)(p21;q23)[20] | ish t(2;11)(p21;q24)(KMT2A-;KMT2A+)[2] | 62 |
| MD16 | 45~46,XY,-3,del(5)(q11.2),add(8)(q24.2),-11,add(14)(q32),-15,-17,+20,+1~2r,+mar[cp20] | nuc ish(KMT2Ax3)[86/200]/(KMT2Ax4)[71/200]/(KMT2Ax5~6)[22/200] | 74 |

**Supplementary Table 3 : Detailed summary of cytogenetic and FISH studies done for the samples prior to our study**.

**qPCR Assay Concordance and Specificity.** The multiplex qPCR assay results were compared against clinical benchmarks (*Supplementary Table 4*). The assay demonstrated high specificity in distinguishing structural fusions from non-fusion abnormalities. Of the samples that tested negative for KMT2A fusions, only **MD13** represented a true wild-type control (normal karyotype, diploid KMT2A). Five samples (**MD1**, **MD3**, **MD4**, **MD9**, **MD10**) harbored KMT2A amplification, and one (**MD16**) displayed monosomy 11 with concurrent KMT2A amplification by FISH. The assay correctly identified all these as negative, confirming its ability to distinguish actionable fusions from non-targetable chromosomal abnormalities.

Sample **MD15** also tested negative; although cytogenetics confirmed a t(2;11) translocation (KMT2A::THADA), this fusion partner is not among the 12 genes included in the current panel design.

| ID | Clinical Classification | Fusion Partner | qPCR Result | Concordance |
| --- | --- | --- | --- | --- |
| MD1 | Negative (Amplification) | WT | Negative | Yes |
| MD3 | Negative (Amplification) | WT | Negative | Yes |
| MD4 | Negative (Amplification) | WT | Negative | Yes |
| MD9 | Negative (Amplification) | WT | Negative | Yes |
| MD10 | Negative (Amplification) | WT | Negative | Yes |
| MD13 | Negative (Wild Type) | WT | Negative | Yes |
| MD16 | Negative (Monosomy 11) | WT | Negative | Yes |
| MD15 | Positive (Rare t(2;11)) | KMT2A::THADA | Negative | No (Target Missing) |
| MD14 | Positive (t(9;11)) | KMT2A::MLLT3 | Positive | Yes |
| MD2 | Positive (t(11;17)) | KMT2A::SEPTIN9 | Positive | Yes |
| MD5 | Positive (t(9;11)) | KMT2A::MLLT3 | Positive | Yes |
| MD6 | Positive (t(9;11)) | KMT2A::MLLT3 | Positive | Yes |
| MD7 | Positive (t(4;11)) | KMT2A::AFF1 | Positive | Yes |
| MD8 | Positive (t(9;11)) | KMT2A::MLLT3 | Positive | Yes |
| MD11 | Positive (t(11;19)) | KMT2A::ELL | Positive | Yes |
| MD12 | Positive (t(11;19)) | KMT2A::MLLT1 | Positive | Yes |

**Supplementary Table 4 Concordance between clinical classification and DIMPLE qPCR results for MD Anderson samples.** Of 16 samples, 15 (93.75%) showed concordant results. The single discordant case (MD15) harbored a rare KMT2A::THADA fusion not included in the DIMPLE panel design.

| 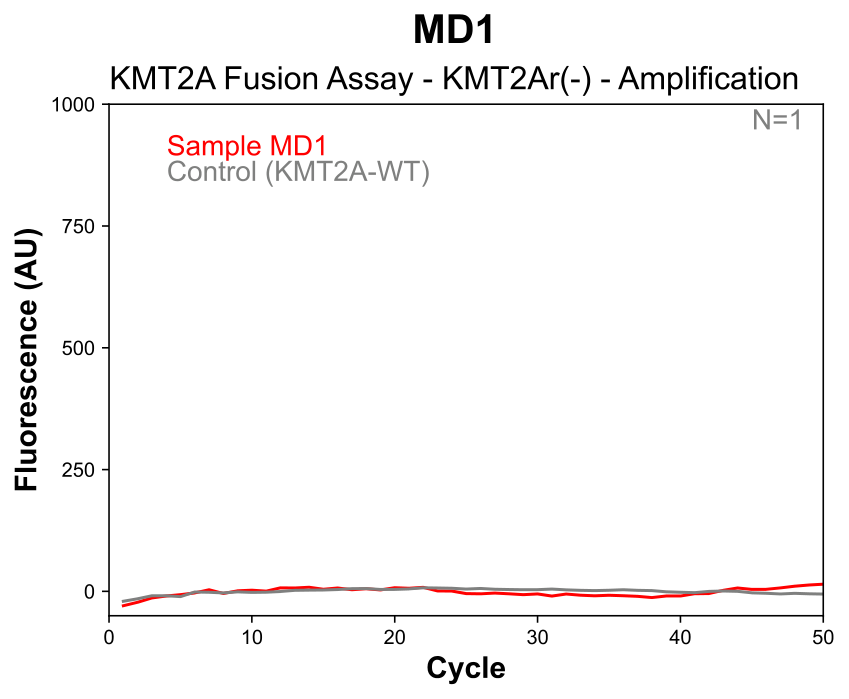 | 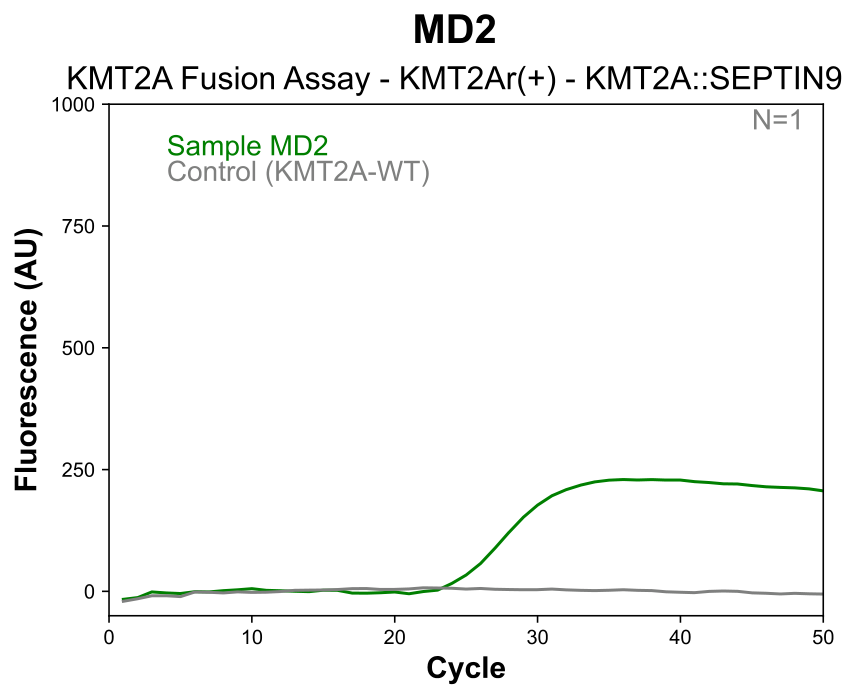 |
| --- | --- |
| 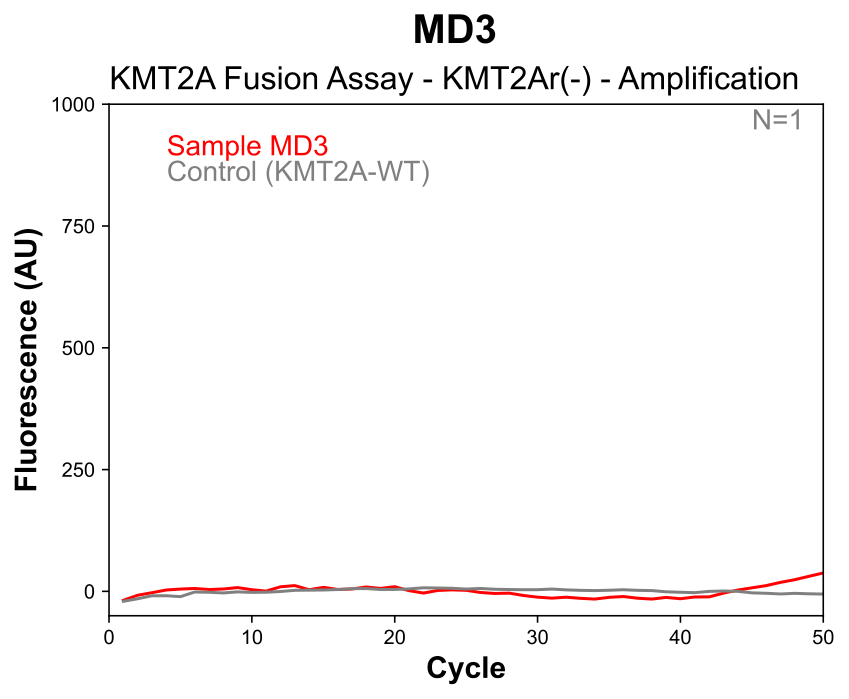 | 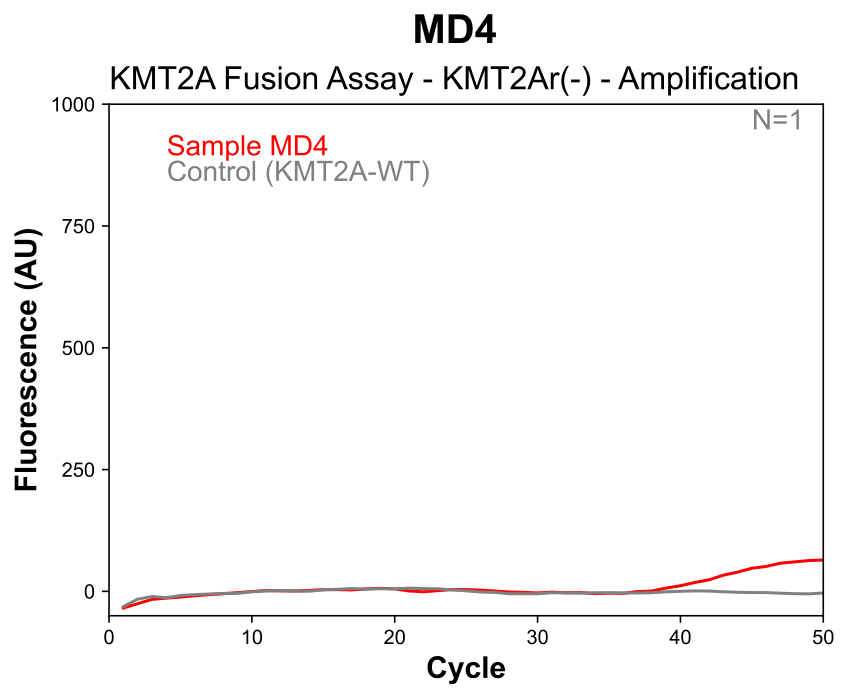 |
| 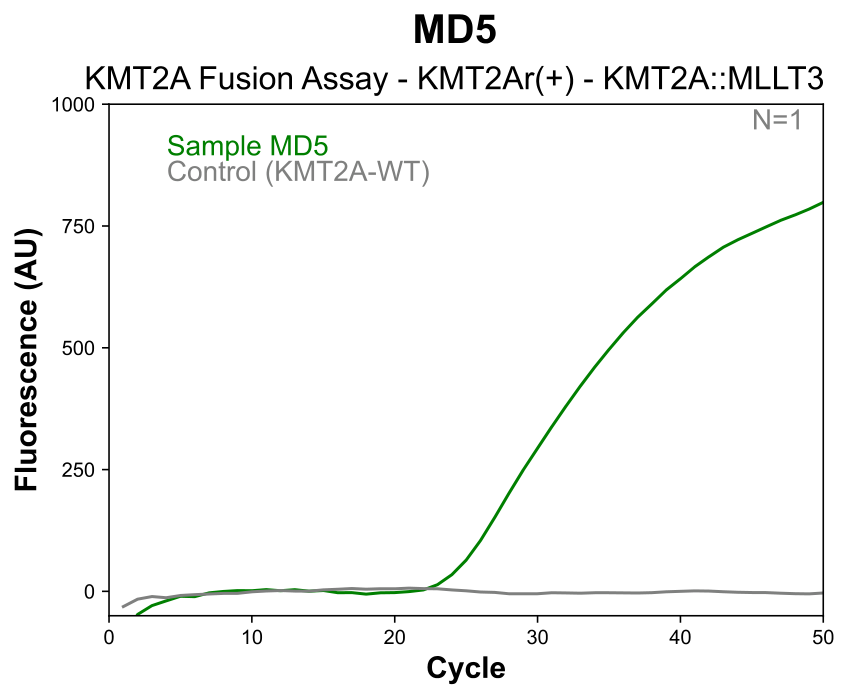 | 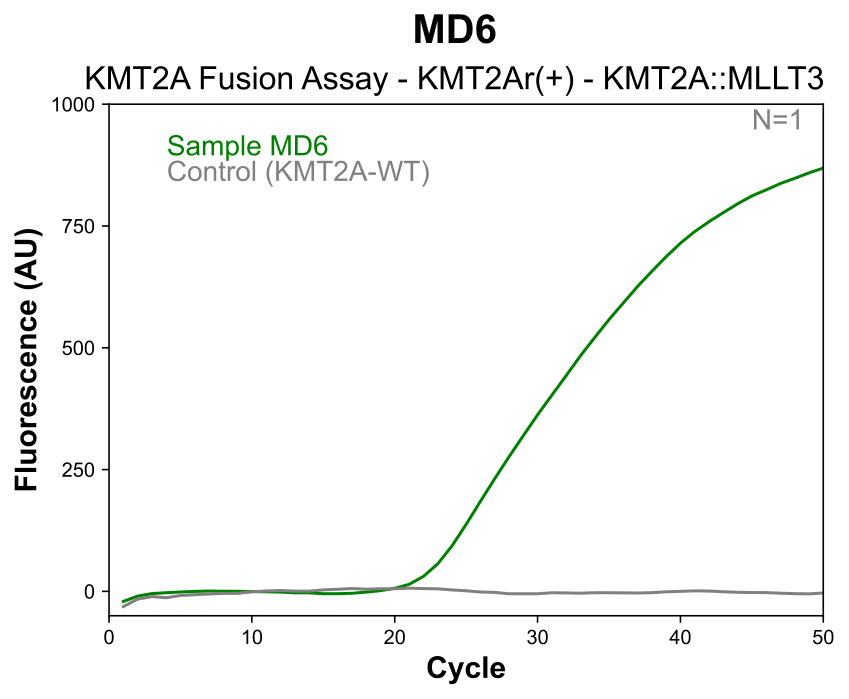 |

| 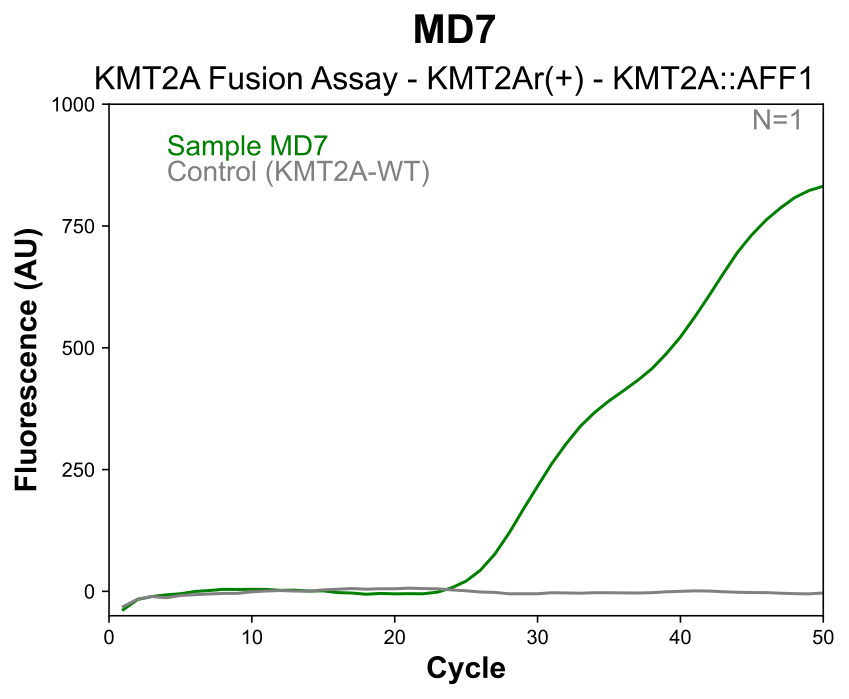 | 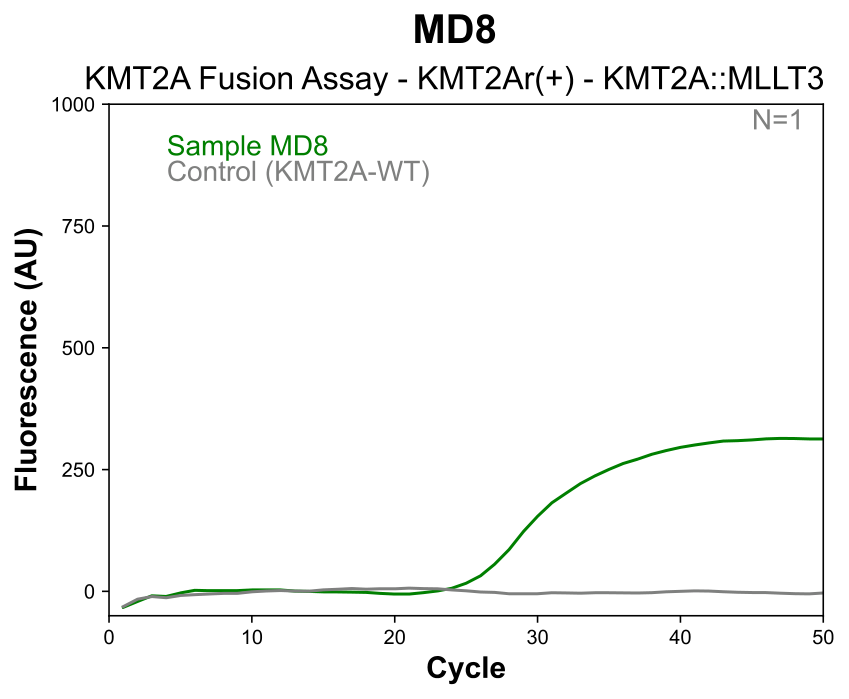 |
| --- | --- |
| 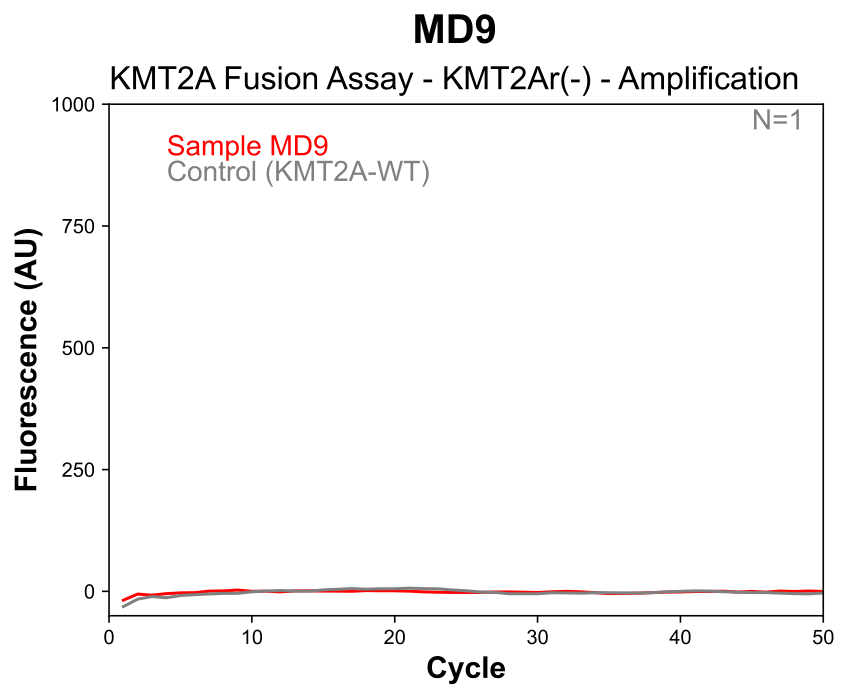 | 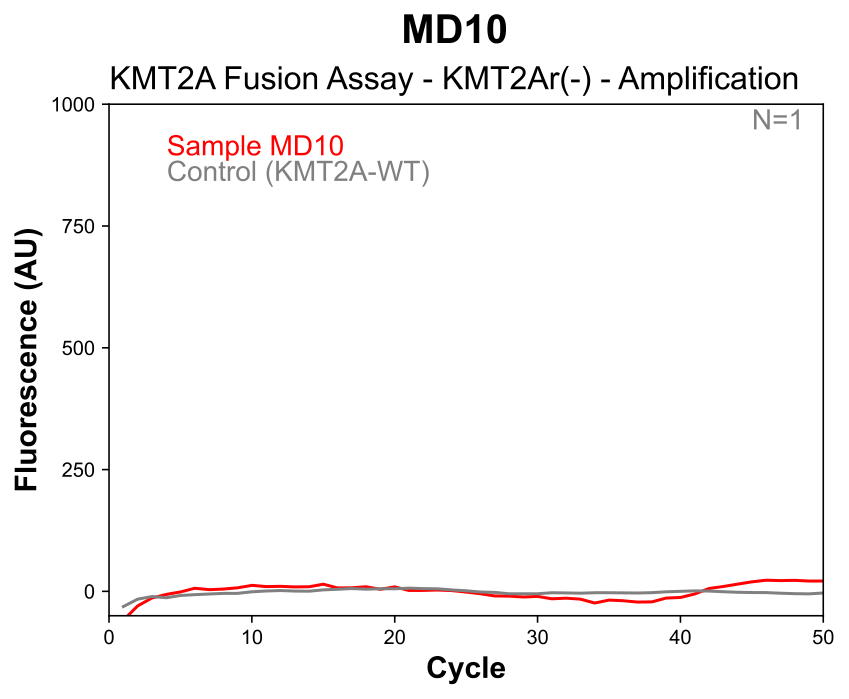 |
| 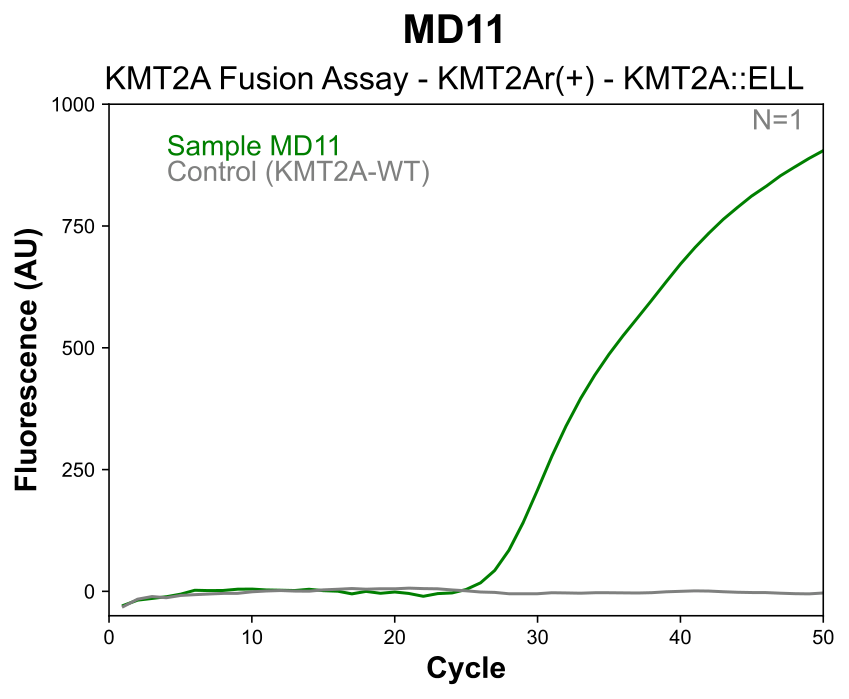 | 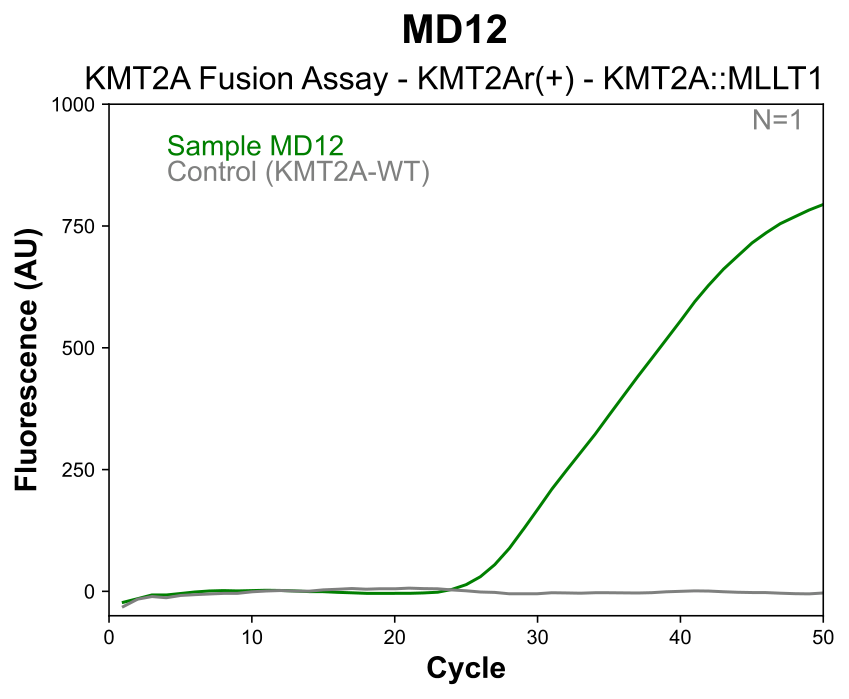 |

| 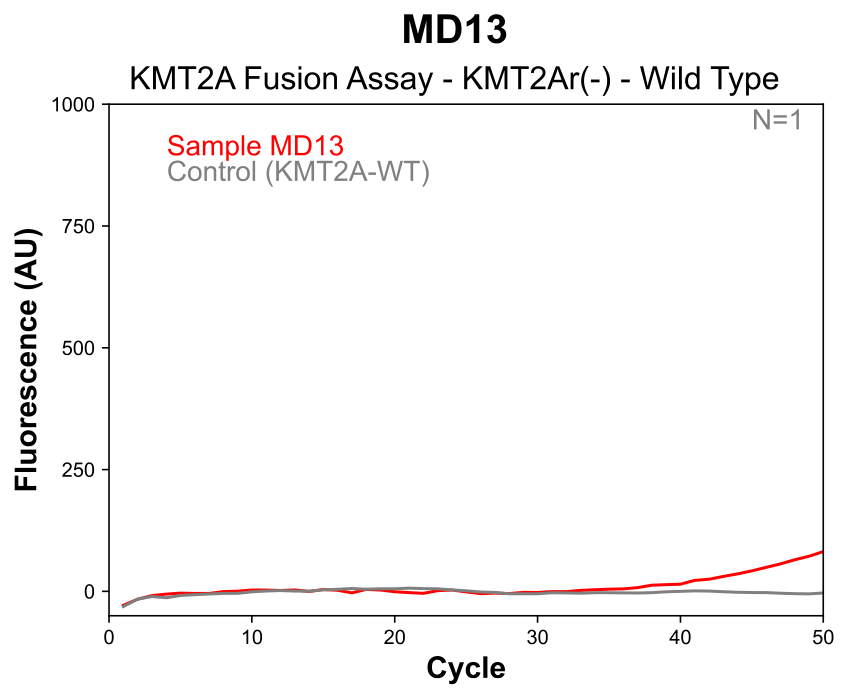 | 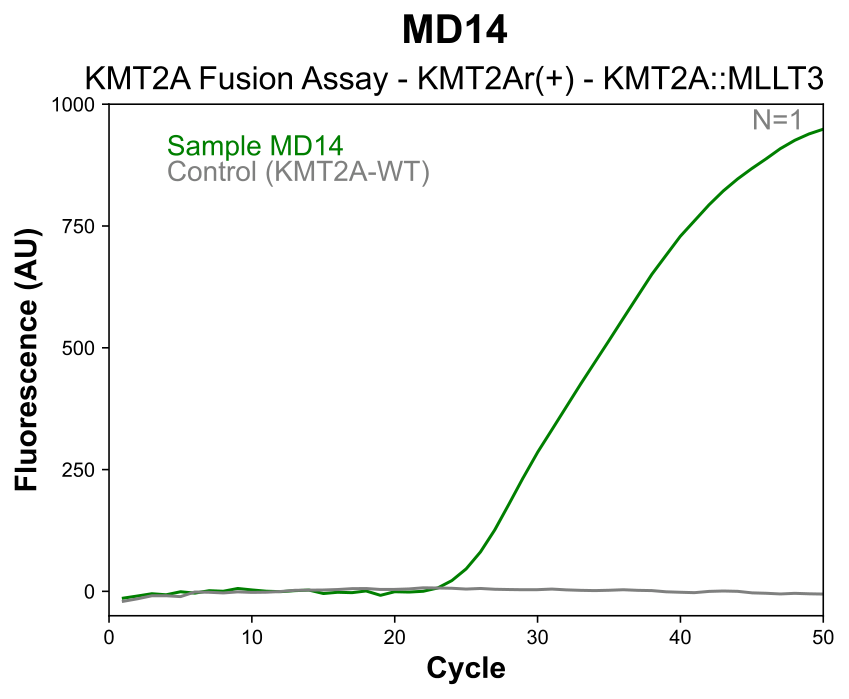 |
| --- | --- |
| 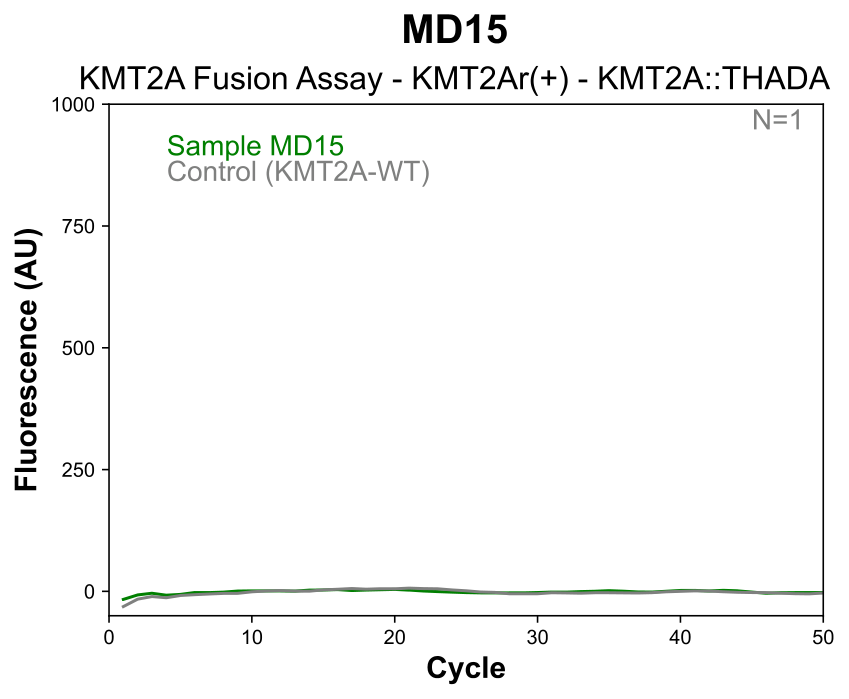 | 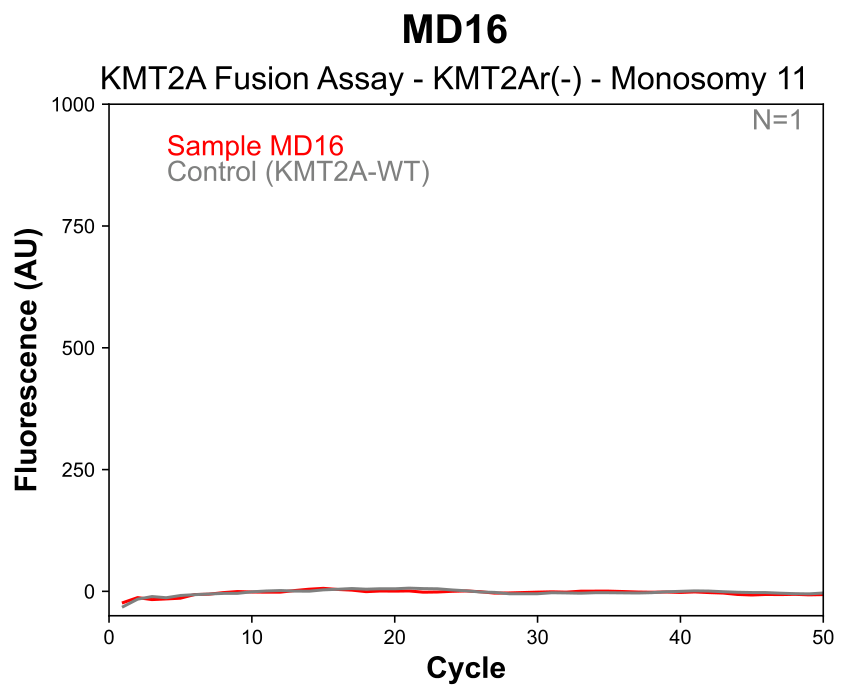 |

**Supplementary Figure 10 qPCR amplification curves for MD Anderson KMT2A samples.** Each panel shows the amplification curve for an individual sample (green: KMT2Ar-positive; red: KMT2Ar-negative) alongside a wild-type negative control included in the assay kit (gray).

**Section S7 | Xinqiao Hospital Assay Results**

Following analytical validation, the multiplex KMT2A assay was kitted using a newly synthesized batch of oligonucleotides. The batch underwent rigorous quality control (QC) to ensure performance consistency before being shipped to Xinqiao Hospital (Chongqing, China) to evaluate assay transferability. The kit was applied to a retrospective cohort of 22 clinical samples previously characterized by Next-Generation Sequencing (NGS). The cohort consisted of 17 samples with confirmed KMT2A rearrangements (KMT2Ar) and 5 sample from. Healty donors - wild-type (WT) for KMT2A.

| 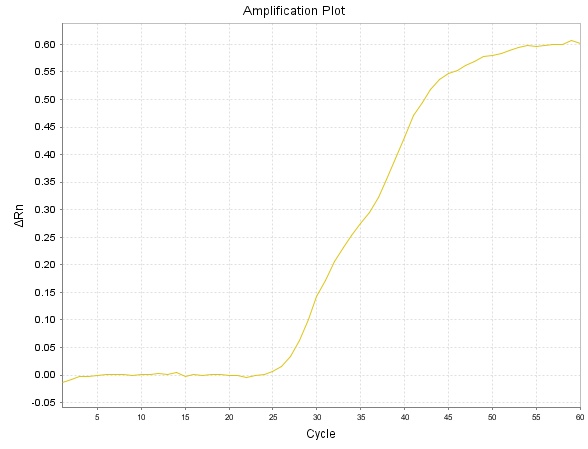  XH1 Sample KMT2A::ELL | 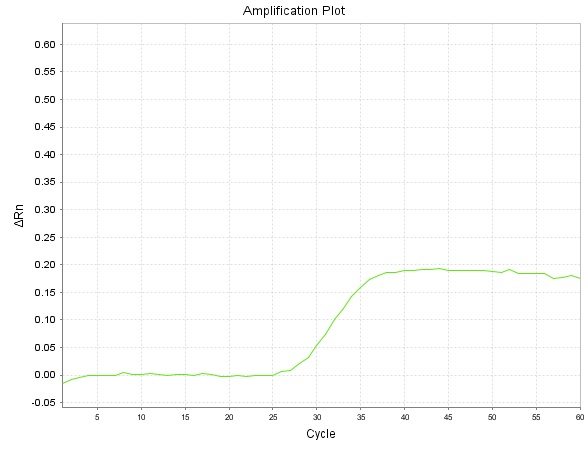  XH2 Sample KMT2A::MLL1 |
| --- | --- |
| 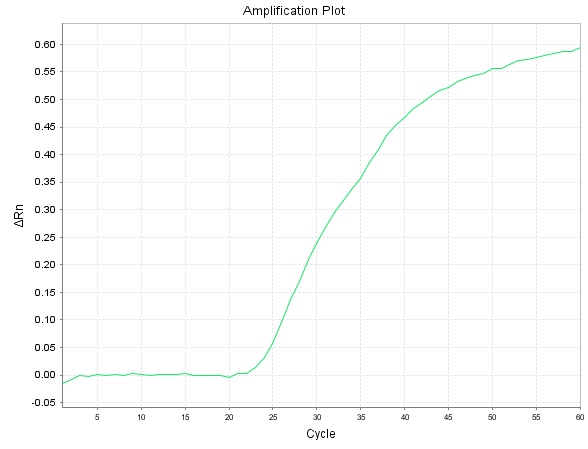  XH3 Sample KMT2A::MLL3 | 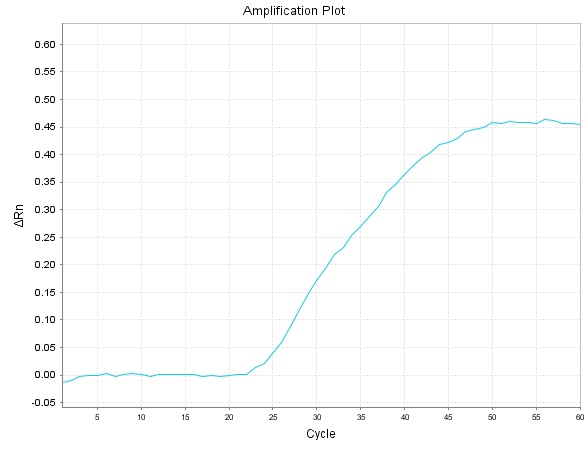  XH4 Sample KMT2A::USP2 |
| 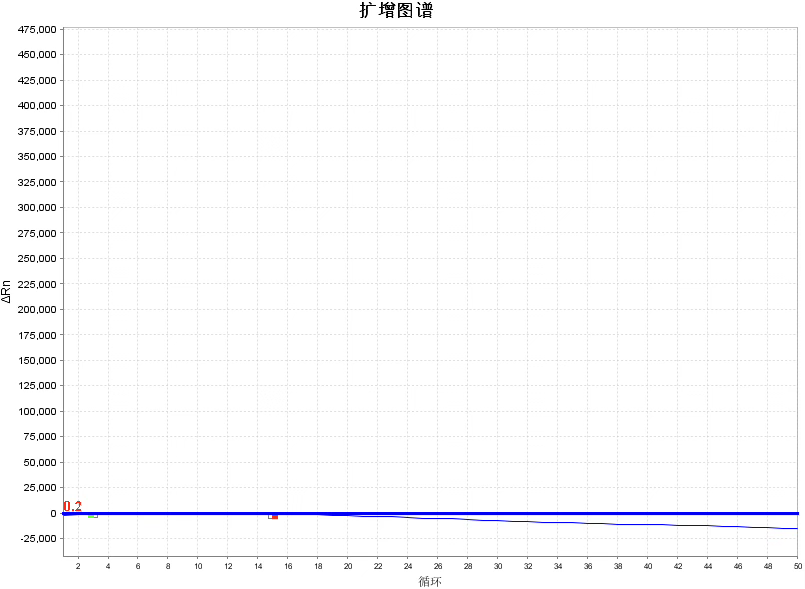  XH5 Sample KMT2A WT | 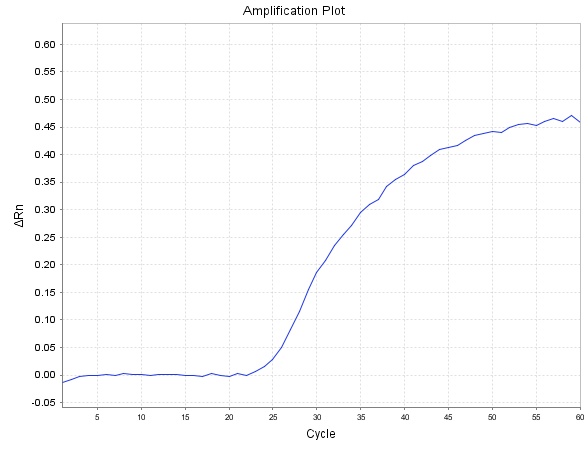  XH6 Sample KMT2A::MLLT3 |
| 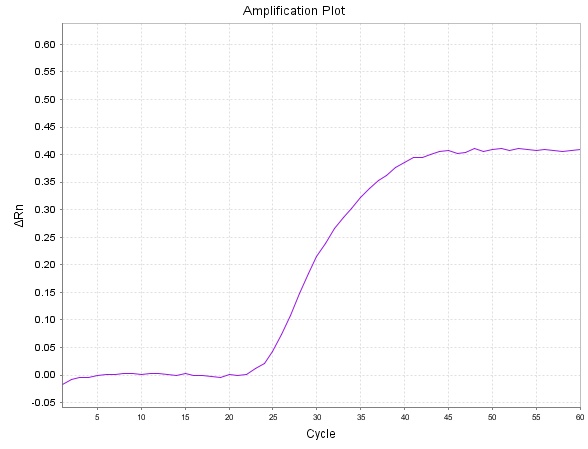  XH7 Sample KMT2A::MLLT3 | 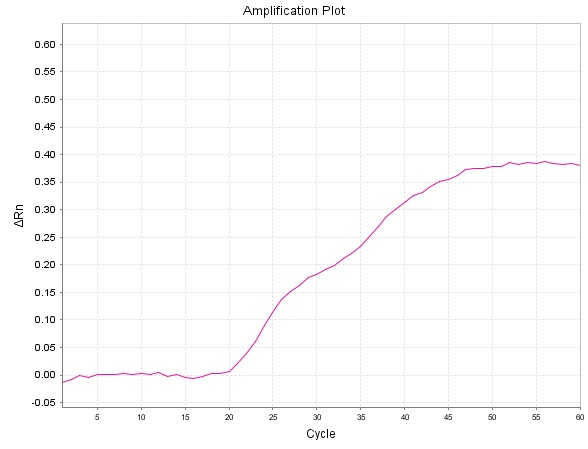  XH8 Sample KMT2A::MLLT4 |
| 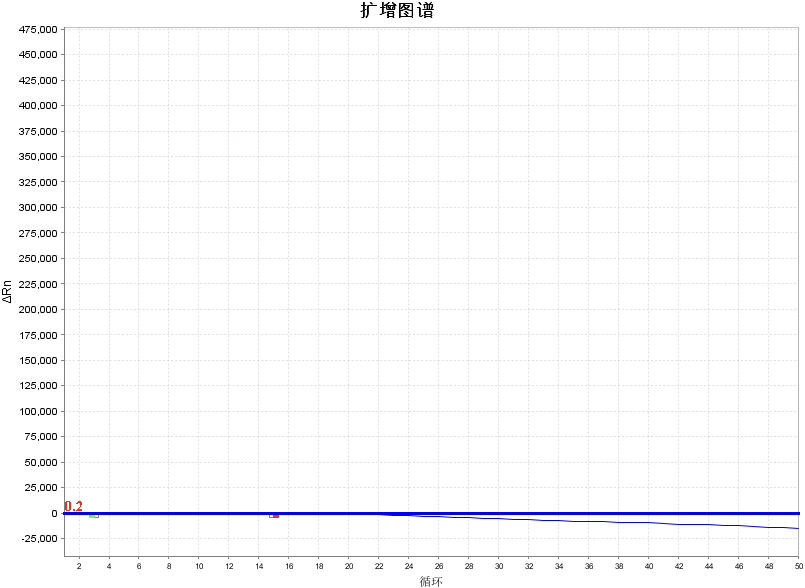  XH9 Sample KMT2A WT | 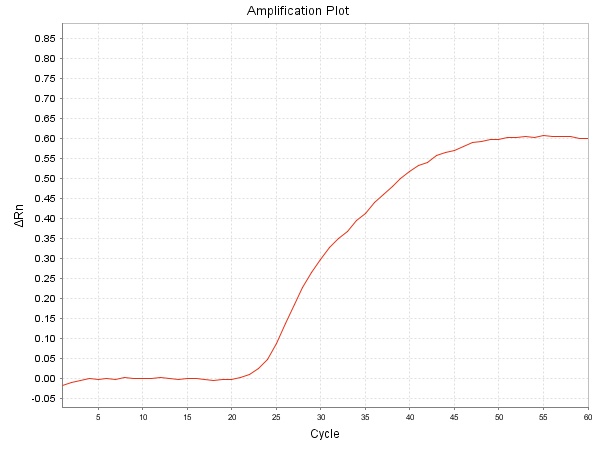  XH10 Sample KMT2A::AFF1 |
| 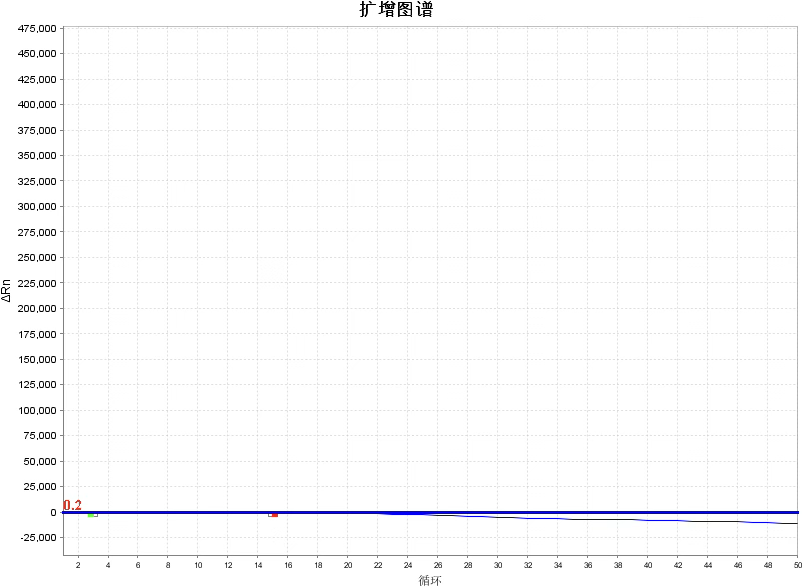  XH11 Sample KMT2A WT | 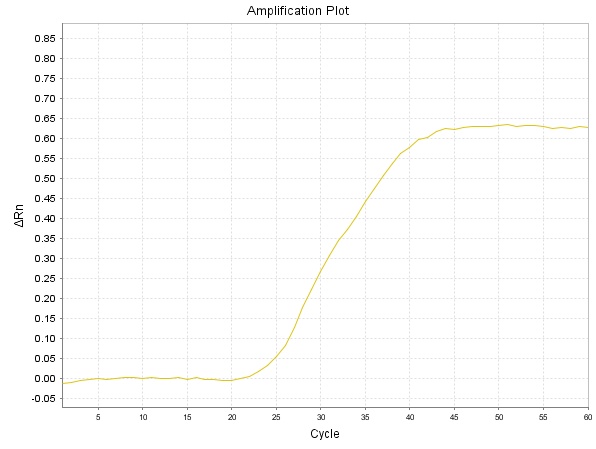XH12 Sample KMT2A::EPS15 |
| 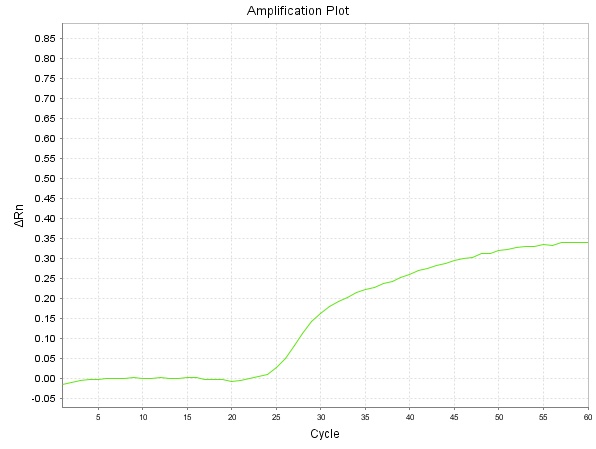  XH13 Sample KMT2A::MLL10 | 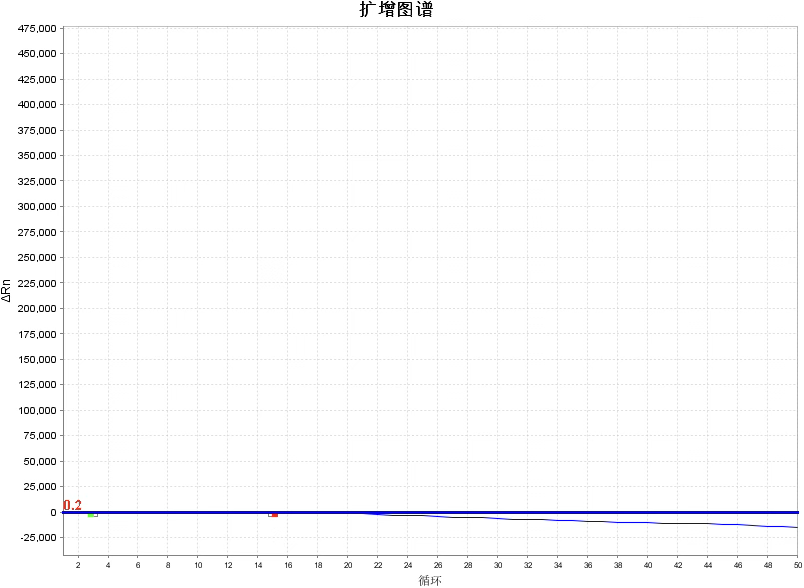  XH14 Sample KMT2A WT |
| 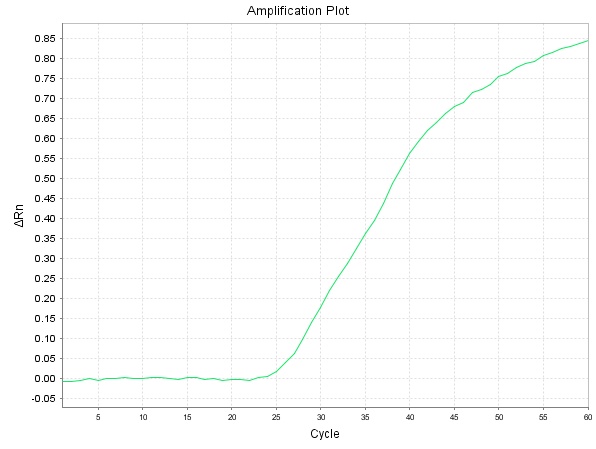  XH15 Sample KMT2A::SEPT6 | 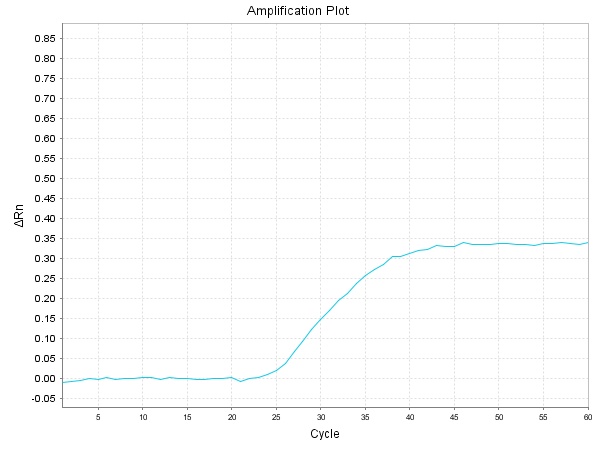  XH16 Sample KMT2A::MLLT6 |
| 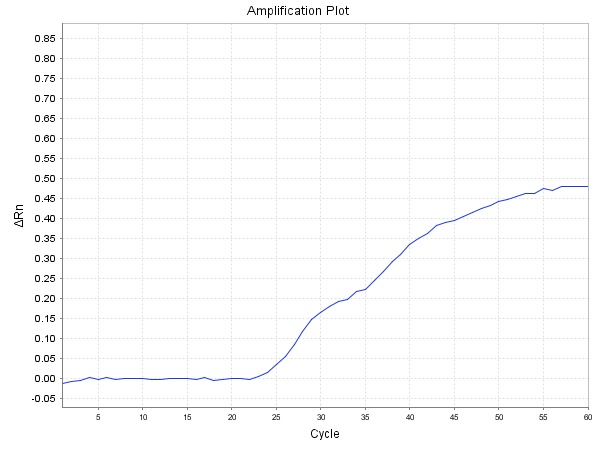  XH17 Sample KMT2A::MLLT11 | 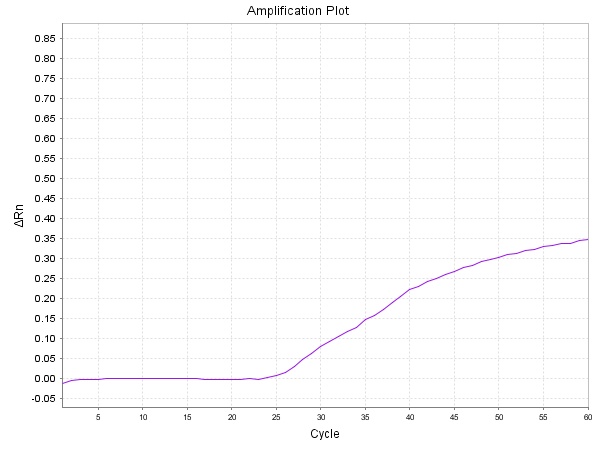  XH18 Sample KMT2A::MLLT11 |
| 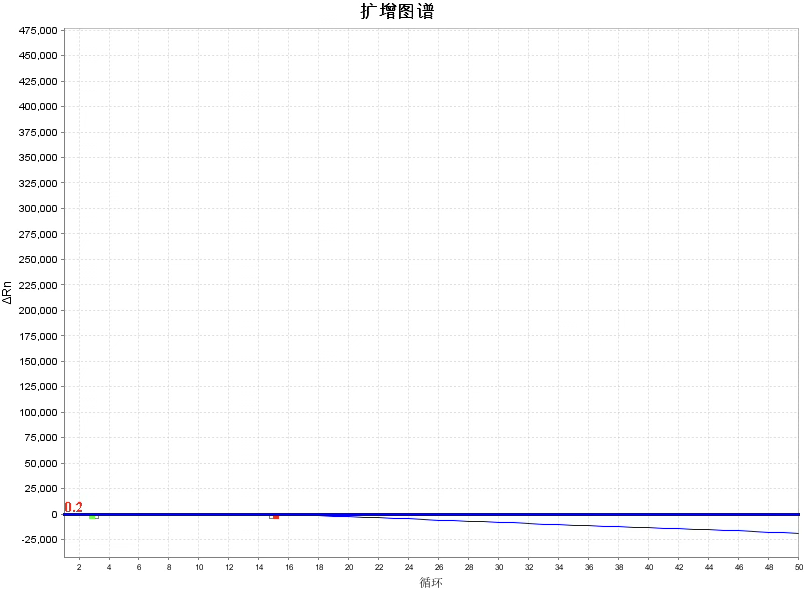  XH19 Sample KMT2A WT | 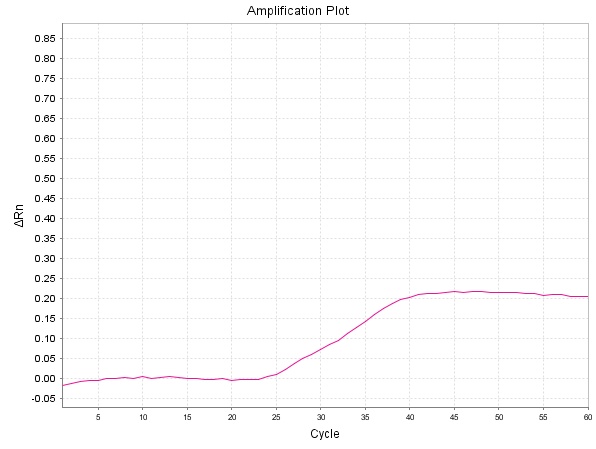  XH20 Sample KMT2A::SEPT6 |
| 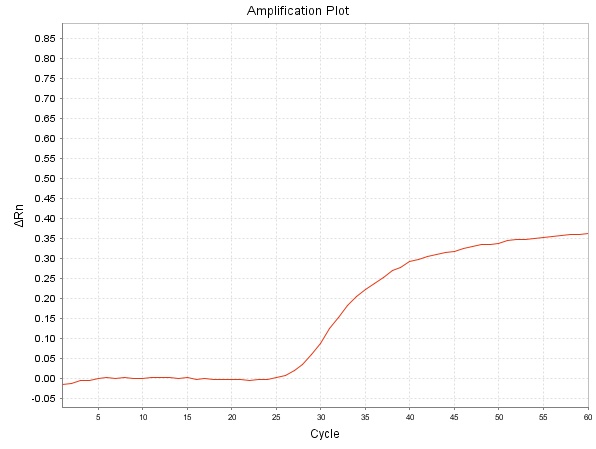  XH20 Sample KMT2A::SEPT9 |  |

Supplementary Figure 11 qPCR amplification curves for Xinqiao Hospital samples. Each panel shows the amplification curve for an individual sample.

1. *Acebedo A, et al. Collaborating across sectors in service of open science, precision oncology, and patients: an overview of the AACR Project GENIE (Genomics Evidence Neoplasia Information Exchange) Biopharma Collaborative (BPC). ESMO Real-World Data and Digital Oncology. 2025;7:100097. doi:10.1016/j.esmorw.2024.100097.*
